# Volumetric Cross-Amplitude Modulation (VxAM) for Transcranial Gas Vesicle Imaging

**DOI:** 10.64898/2026.11.09.765388

**Authors:** Andrew S. Weitz, Hatim Belgharbi, Masashi Sode, Rebecca M. Jones-Sinnott, Rohit Nayak, Trevor Fochtman, Mikhail G. Shapiro, Paul A. Dayton, Gianmarco F. Pinton

## Abstract

Cross-amplitude modulation (xAM) is a commonly used pulse sequence for non-destructive, quasi-artifact-free detection of gas-vesicle (GV) contrast agents, but it has so far been limited to two-dimensional cross-sections acquired with one-dimensional linear arrays. We present the first implementation of xAM on a fully sampled two-dimensional matrix array, extending the half-aperture split of conventional xAM to a family of 2D sub-aperture partitions on a 32*×* 32-element Vermon probe (7.81 MHz, 300 *µ*m element pitch). Angular spectrum simulations compared three primary bisections (X-axis, Y-axis, and diagonal (*D*_1_, *D*_2_)), and the selected Y-split was tested experimentally in GV-loaded tissue-mimicking phantoms. We also simulated an electronic asymmetric-angle sweep as an alternative to the robotic translation used in the experiments, and measured contrast through an *ex vivo* mouse skull. The non-emissive regions between the elevation panels of the Vermon probe accounted for most of the simulated differences among partitions, through both the continuity of the aperture parallel to the bisector plane and the side-lobe structure. At a 6° steering angle, the Y-split gave a contrast-to-tissue ratio (CTR) of *≈* 29 dB for single frames and *≈* 49 dB with 100-frame averaging without the skull, and a contrast-to-noise ratio (CNR) of *≈* 8 dB for single frames and *≈* 12 dB with 100-frame averaging through the skull, where the inclusion remained resolvable. These results give design rules for xAM on matrix arrays. The through-skull result comes from a single *ex vivo* specimen, and *in vivo* work is the next step toward non-invasive, longitudinal, three-dimensional monitoring of GV-labeled glioblastoma and other targets of nonlinear contrast agents.

## I. Introduction

**G**AS vesicles (GVs) are gas-filled protein nanostructures, typically 45–250 nm wide and 100–600 nm long, that form naturally in some bacteria and archaea [1], [2]. Their gas core creates a large acoustic impedance mismatch with the surrounding tissue or blood, so GVs scatter ultrasound efficiently even though they are far smaller than the acoustic wavelength.

GVs differ from linear tissue scatterers in how their acoustic response depends on pressure. At low incident pressure they scatter linearly. Above a buckling threshold *p*_*b*_, which depends on GV type and frequency, the protein shell buckles reversibly and the scattered pressure *P*_*s*_ becomes a nonlinear function of the incident pressure *P*_*inc*_ (**Fig. 1A**) [4], [5]. At higher pressures the shell collapses irreversibly. GVs can also be expressed from acoustic reporter genes (ARGs) in bacteria [6] and mammalian cells [7], and engineered ARG variants give stronger nonlinear contrast [8], so GV contrast can report gene expression *in vivo*.

**Fig. 1.**
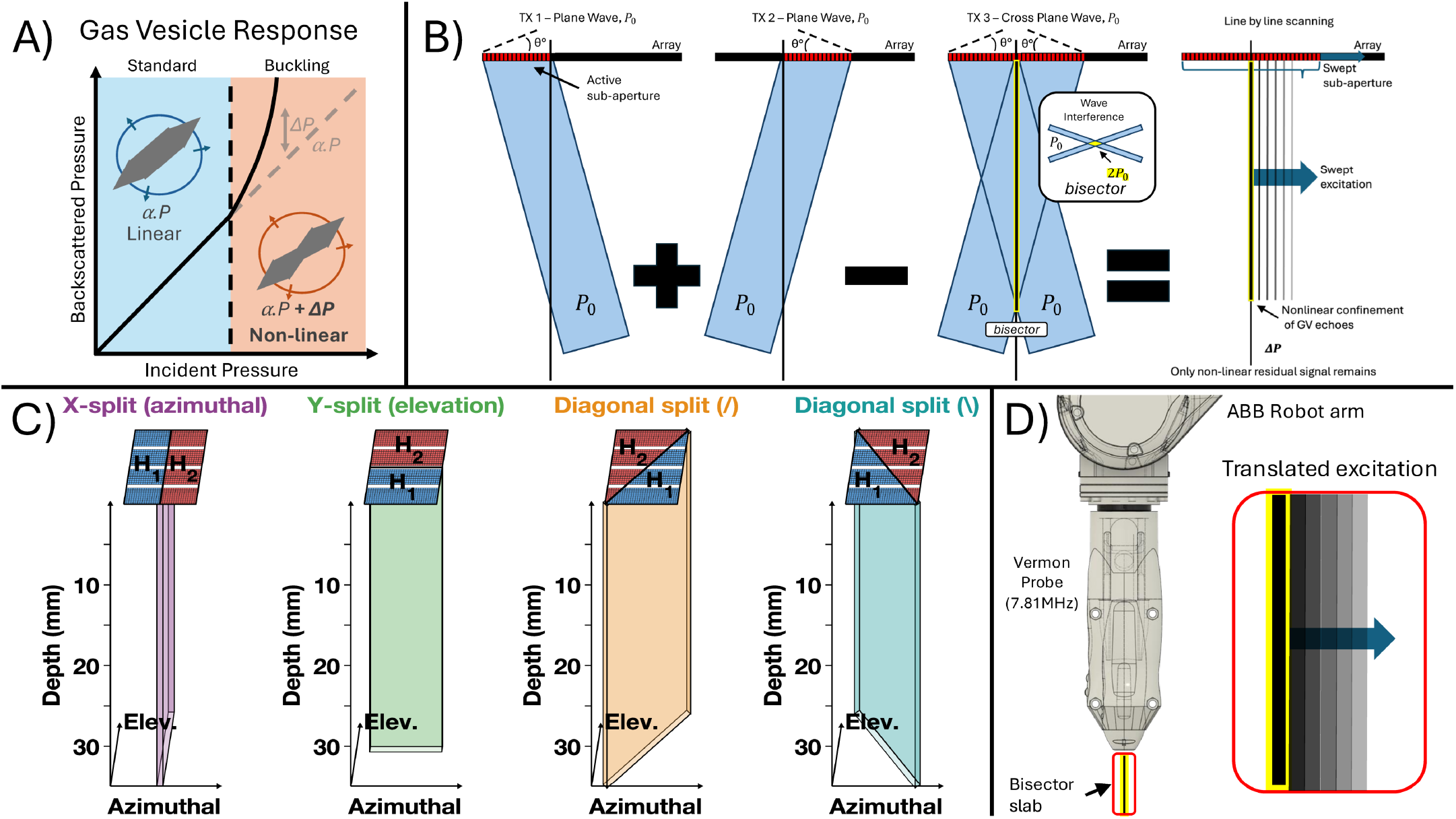
Overview of volumetric cross-amplitude modulation (VxAM). **(A)** Pressure-dependent acoustic response of gas vesicles (GVs): scattered pressure *P*_*s*_ versus incident pressure *P*_*inc*_, with the linear regime below the buckling threshold and the nonlinear increment (Δ*P*) above it. **(B)** The three-event xAM pulse sequence: two half-aperture plane waves *p*_*A*_ and *p*_*B*_, steered symmetrically at +*θ* and *−θ* about the array normal, transmitted individually and then together as a full aperture, with the residual *r*_*xAM*_ = *s*(*p*_*A*_ + *p*_*B*_) *−s*(*p*_*A*_) *−s*(*p*_*B*_) (Eq. 2) and the beams crossing at an angle 2*θ*. Panels (A) and (B) are reproduced from Favre and Maresca [3] with permission of the authors. **(C)** Aperture-partitioning strategies on the fully sampled 2D matrix array, shown as simulated bisecting-beam fields: X-split (azimuthal bisection), Y-split (elevation bisection), and the two diagonal splits *D*_1_ and *D*_2_, split along axes at +45° and *−*45°. **(D)** Volumetric acquisition by robotic translation: the Vermon 7.81 MHz matrix probe forms a fixed xAM bisector slab that an ABB robot arm sweeps through the volume in elevation to synthesize the 3D VxAM dataset.

Imaging GVs with high specificity requires separating their *local* nonlinear scattering from the *cumulative* nonlinear distortion that a wave acquires as it propagates through tissue. In a medium with coefficient of nonlinearity *β* = 1 + *B*/2*A*, a finite-amplitude wave generates harmonics and steepens progressively along its path [9]. For a plane wave of amplitude *p*_0_ propagating a distance *z*, the amplitude of the cumulative second harmonic scales as

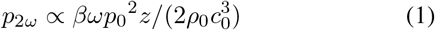

where *ω* is the angular frequency, *ρ*_0_ the ambient density, and *c*_0_ the sound speed. This propagation artifact arises with no nonlinear scatterer present and grows linearly with distance. Amplitude modulation (AM) transmits collinear beams at half and full amplitude and forms the residual *s*(*p*_0_) *−*2 *s*(*p*_0_*/*2), where *s(p)* is the received backscattered signal from a transmission at peak pressure *p*. This residual vanishes for a linear scatterer, but the cumulative nonlinearity of Eq. 1 is quadratic in *p*_0_ and leaves a nonzero residual from tissue alone, which limits specificity [4]. The size of this residual depends strongly on the shape of the transmitted wavefronts [10].

Building on the detection of contrast agents with interacting, quasi-counter-propagating wavefronts [11], cross-amplitude modulation (xAM), introduced by Maresca *et al*., addresses this problem by changing the geometry of the wave interaction [4]. Instead of collinear beams that accumulate nonlinearity along a shared axis, xAM splits an active sub-aperture of the linear array into two halves and uses three transmit events: the two halves each transmit a plane wave steered at +*θ* and *−θ* from the array normal, each fired alone, and a third event fires both halves together (**Fig. 1B**). The xAM residual is

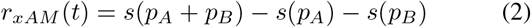

where *p*_*A*_ and *p*_*B*_ are the pressure fields from sub-apertures *A* and *B*. Steering the two beams so that they cross at angle 2*θ* confines their nonlinear interaction to a narrow region along the bisector [4]. Away from the bisector the beams do not overlap. Each propagates independently, the nonlinearity accumulated in the joint transmission equals the sum of that accumulated in the two single transmissions, and Eq. 2 returns zero. Because cumulative nonlinearity is quadratic in pressure **(Eq. 1)**, the only propagation term that can survive the subtraction is the cross-term *∝p*_*A*_*p*_*B*_, which is generated only where the beams overlap, the only region in which their nonlinear interaction source terms are nonzero [12].

Local nonlinear scattering from GVs, on the other hand, survives the subtraction. On the bisector the incident pressure is *p*_*A*_ + *p*_*B*_. If this exceeds the GV buckling threshold *p*_*b*_ while neither *p*_*A*_ nor *p*_*B*_ alone does, then *s*(*p*_*A*_ + *p*_*B*_) contains nonlinear GV scattering and *s*(*p*_*A*_) and *s*(*p*_*B*_) do not, and the resulting nonzero *r*_*xAM*_ arises from the contrast agent. Each transmit triplet yields one image line along its bisector, and stepping the active sub-aperture element by element across the 1D array builds up a 2D xAM image [4]. The cancellation is not complete, however. The propagation cross-term also survives on the bisector and leaves a residual propagation arti-fact, most visible distal to strongly scattering inclusions. This artifact decreases as the steering angle *θ* increases and nears extinction above *θ ≈* 16.5° [4], and in simulation it is one to two orders of magnitude smaller than the propagation residual of AM with focused, plane, or diverging wavefronts [10]. Because it suppresses cumulative propagation nonlinearity to this small residual while keeping local scattering nonlinearity, xAM has become a commonly used pulse sequence for non-destructive, quasi-artifact-free GV imaging [3], [4].

Related amplitude-modulation methods have since been developed for ultrafast plane-wave imaging (uAM) [13], harmonic detection (HxAM) [14], singular-value-decomposition filtering (AM-SVD) [15], [16], and synthetic nanobubble contrast agents [17]. Favre and Maresca derived analytical expressions for the X-wave beam width *w(z)* and the distance *d*_*SL*_ between primary and secondary lobes as functions of steering angle and wavelength [3].

All of these implementations image two-dimensional cross-sections with one-dimensional linear arrays, whereas tumor angiogenesis, immune cell trafficking, and bacterial colonization unfold in three dimensions. In nonlinear sound-sheet microscopy (NSSM), Heiles *et al*. extended xAM to 3D with a high-frequency row–column-addressed (RCA) array, whose orthogonal row and column elements transmit cross-propagating non-diffracting beams that intersect along a thin acoustic sheet. Translating the sheet through the volume yields a stack of 2D xAM slices, and NSSM imaged tumor gene expression with genetically encoded GVs and the cerebral microvasculature in 3D [18]. The RCA geometry, however, fixes the cross-propagation to the row-column bisector plane, which rules out multi-directional angular sampling, and it bounds the usable depth by the cross-propagation depth (*∝* cot *θ*), so increasing *θ* to suppress propagation artifacts also shortens the sound sheet.

Separately, Lee *et al*. introduced “Takoyaki” ultrasound, which uses a multiplexed matrix array to scan multiple focal points through a 3D volume simultaneously, and achieved realtime volumetric contrast imaging of GVs and ARG-expressing glioblastoma tumors [19]. Takoyaki shows that a matrix array can detect GVs volumetrically, but it uses focused transmissions instead of cross-propagating plane waves. Propagation along each focused scanline is collinear, so Takoyaki does not inherit the propagation-artifact suppression of xAM. On the same paneled array, driven through a 4-to-1 multiplexer in which elements at equivalent positions in the four banks share delays, Lee *et al*. also implemented a volumetric xAM sequence and a cylindrically focused sheet AM sequence (Sheet-pAM). Their xAM split the aperture along the lateral axis, giving a bisector perpendicular to the inter-panel gaps, and its contrast was strongly anisotropic. The orthogonal orientation was not tested, because the gaps were expected to disrupt the crossing waves and the multiplexer restricted the aperture [19].

Here we present the first implementation of volumetric cross-amplitude modulation (VxAM) on a fully sampled two-dimensional matrix array. With *N× N* independently addressable elements in both the lateral (*x*) and elevation (*y*) dimensions, a matrix array can divide its aperture into two halves in many more ways than a 1D array can. We compare three primary partitions with angular spectrum method (ASM) simulations and test the selected one experimentally in tissuemimicking phantoms **(Fig. 1C)**. The three are (i) X-axis bisection, which reproduces conventional 1D xAM in the lateral-axial plane, (ii) Y-axis bisection, which rotates the cross-propagation plane into the elevation-axial plane, and (iii) diagonal bisection, which orients the cross-propagation plane at 45°/ *−*45° to the array axes. The partitions trade off field uniformity, peak-positive pressure (PPP) at the beam intersection, and artifact suppression, which we characterize with simulations in homogeneous and aberrating media and with phantom experiments.

A VxAM transmission produces a volumetric slab at the beam intersection, so imaging a larger volume requires sweeping or stacking that slab. We compare two ways of doing so, which differ in acquisition speed, field-of-view extent, and implementation complexity: *native* VxAM, which steers the slab electronically with asymmetric angles from a single fixed probe position, and *translated* VxAM, which moves the probe in elevation with a robot arm **(Fig. 1D)**. Both are described in Sec. II-I.

## II. Methods

### A. Matrix Array Transducer and Synchronized Multi-Vantage Acquisition

All data were acquired with a fully addressed 1024-element two-dimensional matrix array (Vermon, Tours, France) with center frequency *f*_0_ = 7.81 MHz (*−*6 dB fractional bandwidth *≈*60%; wavelength *λ*_0_ *≈* 197 *µ*m at *c*_0_ = 1540 m/s). Table I lists the acquisition and simulation parameters used throughout. The 32 *×*32 elements are arranged as four 32 *×*8 subpanels (32 azimuthal*×* 8 elevation elements) stacked along the elevation axis and separated by non-emissive regions. This *disjointed* layout differs from that of a monolithic matrix **(Fig. 2)**. These inter-panel non-emissive regions add grating lobes and spatially modulate the effective element response. A uniform-pitch approximation captures neither effect, so both beam synthesis and simulation must model the non-emissive regions explicitly.

**Table I.**
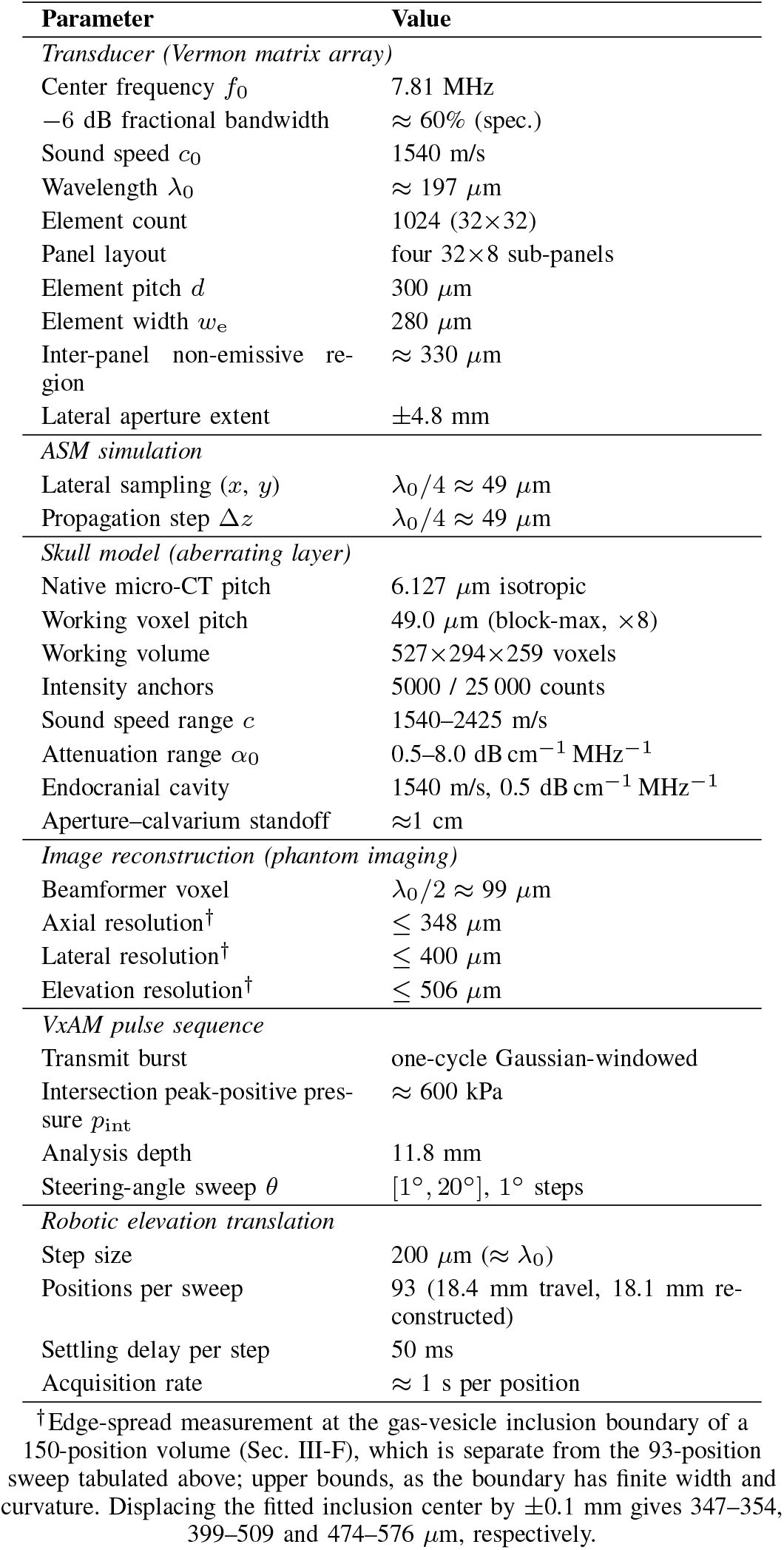
Acquisition and Simulation Parameters Used Throughout This Work.

**Fig. 2.**
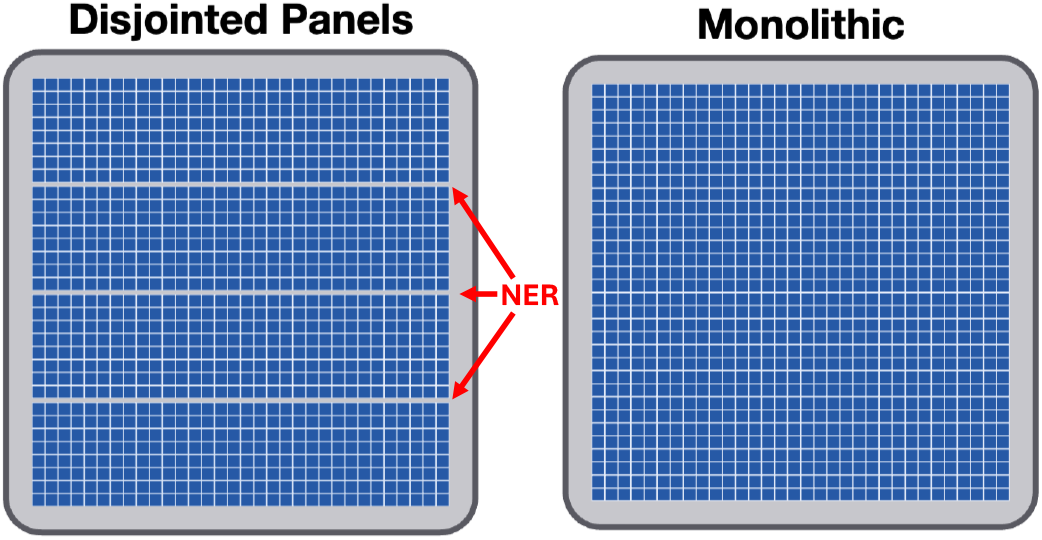
Disjointed (paneled) versus monolithic matrix-array topology. The Vermon 32 *×*32 aperture is built from four 32*×* 8 sub-panels stacked along the elevation axis and separated by non-emissive regions (NER), in contrast to an idealized monolithic matrix of uniform pitch.

To address all 1024 channels, four Verasonics Vantage 256 research scanners (Verasonics, Kirkland, WA) were operated together, as previously demonstrated with this array [20]. The scanners were connected to the probe through a four-way transducer adapter and synchronized in a primary–secondary configuration with a shared sampling clock and transmit trigger. This gives fully programmable element-wise transmit delays over the whole aperture, which xAM needs for arbitrary sub-aperture partitions and for firing both sub-apertures simultaneously.

### B. Aperture Partitioning Strategies

On a 2D matrix, sub-apertures *A* and *B* can be any two non-overlapping element sets that together tile the active aperture. Elements are indexed by (*i, j*), with *i* the azimuthal (*x*) row index and *j* the elevation (*y*) column index, each running from 0 to 31.

#### 1) X-axis bisection (X-split)

The aperture is divided along the azimuthal (*x*) axis into two 16*×* 32 half-apertures, *A*_*x*_ =*{* (*i, j*) : *i ≤*15*}* and *B*_*x*_ = *{* (*i, j*) : *i≥* 16 *}*. Each half transmits a plane wave steered by *± θ* in the *xz* plane, reproducing conventional 1D xAM in the lateral-axial plane **(Fig. 3, X-split)**.

**Fig. 3.**
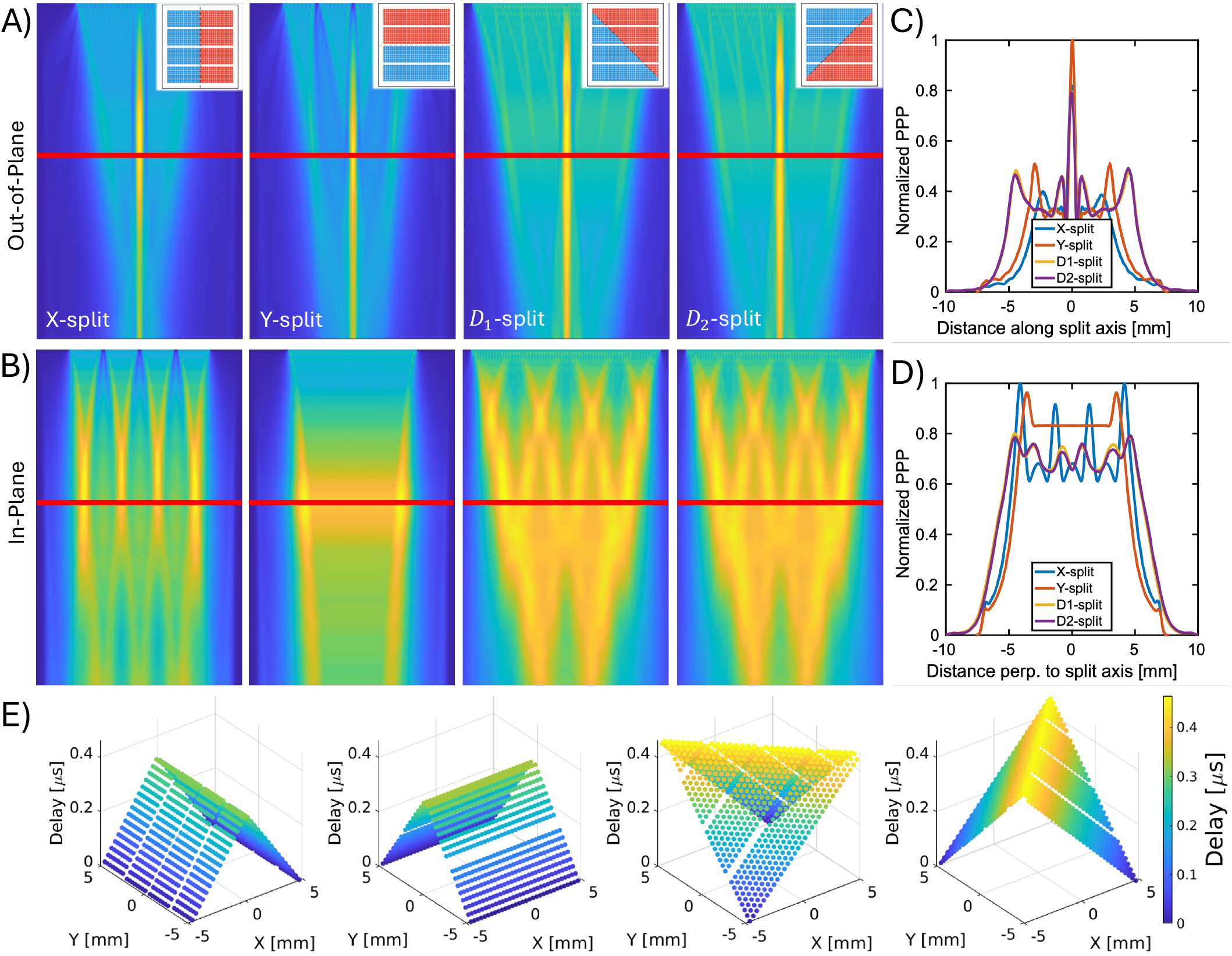
Sub-aperture partitions, their simulated bisecting-beam fields, and their transmit-delay profiles. Profiles are taken in two directions relative to the bisector plane throughout: *out-of-plane* (along the split axis, perpendicular to the bisector) and *in-plane* (perpendicular to the split axis, parallel to the bisector). **(A)** Out-of-plane normalized pressure fields for the X-, Y-, *D*_1_-(split axis +45°), and *D*_2_-(split axis *−*45°) splits, with the half-aperture partition inset in each; the red line marks the 11.8 mm analysis depth. **(B)** Corresponding in-plane pressure fields. **(C)** Normalized peak-positive-pressure (PPP) out-of-plane profiles at 11.8 mm, showing the main lobe and side lobes of each partition. **(D)** Normalized PPP in-plane profiles. **(E)** Element-wise transmit-delay maps (in *µ*s) across the 32*×*32 aperture. In every partition the two sub-apertures are steered to *±θ* so that their beams cross symmetrically at the bisector.

#### 2) Y-axis bisection (Y-split)

The aperture is divided along the elevation (*y*) axis into two 32*×* 16 half-apertures, *A*_*y*_ =*{* (*i, j*) : *j ≤*15*}* and *B*_*y*_ =*{* (*i, j*) : *j≥* 16*}*, which transmit plane waves steered by*± θ* in the *yz* plane. The cross-propagation plane is then elevation-axial, and the bisector runs parallel to the inter-panel non-emissive regions **(Fig. 3**, **Y-split)**.

#### 3) Diagonal bisection (D_1_, D_2_)

The aperture is divided by a 45°/ *−*45° line through its geometric center, and each element is assigned by the sign of its *physical* position relative to that line: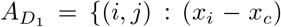 + (*y*_*j*_*− y*_*c*_) *<* 0 *}* for *D*_1_, split along the +45° axis, and 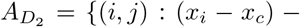 (*y*_*j*_ *−y*_*c*_) *<* 0*}* for *D*_2_, split along the *−*45° axis, with *B* the complement in each case and (*x*_*c*_, *y*_*c*_) the aperture center. Both partitions split the 1024 elements exactly in half, so the two sub-apertures radiate equal amplitude and their beams cross on the array axis. Because the elevation axis carries the three inter-panel non-emissive regions, this geometric partition is *not* the same as the index-space condition *i* + *j ≤* 31 (for *D*_1_), which would assign 528 elements to one sub-aperture and 496 to the other and shift the crossing by *≈* 75 *µ*m. The two steering directions lie in the diagonal planes, so the cross-propagation samples a different region of the (*k*_*x*_,*k*_*y*_) spectrum than the X- or Y-split does **(Fig. 3**, *D*_1_**/***D*_2_ **splits)**.

Each sub-aperture transmitted a plane wave using element-wise delays

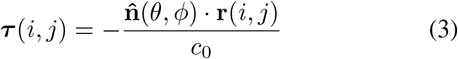

where **r**(*i, j*) is the 3D position of element (*i, j*) on the array, and 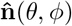 is the unit vector in the desired propagation direction, parameterized by steering angle *θ* (measured from the array normal) and azimuth angle *ϕ*:

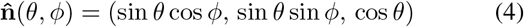

The sound speed was set to *c*_0_ = 1540 m/s. The azimuth *ϕ* sets the split: *ϕ* = 0, *π/*2, *π/*4, and *− π/*4 for the X-, Y-, *D*_1_-, and *D*_2_-splits, respectively. Within each split the two sub-apertures were steered to equal and opposite angles (*± θ*) so that their beam axes crossed symmetrically. Apodization was rectangular within each sub-aperture, with no additional lateral or elevation apodization.

### C. VxAM Pulse Sequence

The three-event VxAM sequence **(Eq. 2)** consisted of (i) sub-aperture *A* alone, (ii) sub-aperture *B* alone, and (iii) sub-apertures *A* and *B* fired simultaneously, all at full amplitude **(Fig. 1B)**. Each transmission was a one-cycle coherent Gaussian-windowed burst at *f*_0_ = 7.81 MHz. On receive, the full 1024-element aperture was active and raw per-channel radio-frequency data were stored, with all receive delays applied in software during reconstruction (Sec. II-J). For angle sweeps, *θ* was varied in 1° increments over [1°, 20°], covering angles both below and above the *θ >* 16.5° at which Maresca *et al*. report that the residual propagation artifact approaches complete extinction [4].

To set the operating amplitude, the transmit pressure was ramped at a fixed probe position on a scatterer-free agarose phantom, so that scattering in the background would not alter the pressure reaching the GV inclusion. The drive voltage was stepped from 1.6 to 14 V in 0.2 V steps (63 levels), corresponding to intersection pressures of *≈* 150–2700 kPa through a voltage–pressure relation measured with a hydrophone over 2–10 V and extrapolated linearly outside that range **(Fig. S1)**. At each level the inclusion was insonified until the reconstructed residual stabilized before the 100-frame batch was acquired, so all reported metrics are steady-state values for the GV population that survives at that amplitude. The operating point was set at the maximum of the annulus-referenced contrast ratio (Sec. III, **Fig. S2**), a PPP at the beam intersection of *p*_int_ *≈* 600 kPa (591 kPa at the 4.0 V calibration node of Fig. S1). This is the coherent sum of the fields of sub-apertures *A* and *B*. Each alone delivers about half that amplitude at the analysis depth and produced no detectable nonlinear GV response. The relation of this operating point to published collapse pressures is discussed in Sec. IV-C.

### D. Angular Spectrum Method Simulation Framework

The forward beam physics of each partition was simulated with a custom in-house ASM solver [21]. ASM decomposes the transmit field at the source plane *z* = 0 into its two-dimensional spatial Fourier spectrum and propagates each plane-wave component forward in *z* with a multiplicative propagator (5):

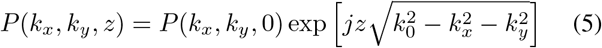

where *P* (*k*_*x*_, *k*_*y*_, 0) = ℱ_*xy*_ *p{*(*x, y*, 0)} is the spatial Fourier transform of the source field, *k*_0_ = 2*πf*_0_*/c*_0_ is the wavenumber, and the square root separates propagating components 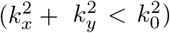 from evanescent ones, which decay exponentially. The field at any depth is recovered by inverse transformation. The propagator of (5) is evaluated at *f*_0_ only, so the simulation is monochromatic. The experimental transmit is a short burst whose measured spectrum spans *≈* 5.6–9.7 MHz at *−*6 dB, a fractional bandwidth of *≈* 53% of *f*_0_ **(Fig. S1)**, within the transducer’s specified *≈* 60%. Finite bandwidth lowers the side-lobe level, shorter pulses giving fewer and weaker lobes [3], whereas the lobe spacing is set by the crossing geometry and matches pulsed hydrophone measurements [3].

The X-wave side lobes are spaced by *d*_*SL*_ = *λ*_0_*/*(2 sin *θ*) in both cases, but at that offset the two pulses arrive one full period apart, so their envelopes overlap by at most *≈* 0.25 and the pulsed side lobes are weaker than a continuous-wave calculation predicts. Simulated side-lobe levels are therefore upper bounds, while the spacing *d*_*SL*_ used as a selection criterion in Sec. II-E is unaffected.

Each simulated transmission was built by applying the element-wise transmit delays across the source plane. The grid was sampled isotropically at *λ*_0_*/*4 *≈* 49 *µ*m in *x, y*, and *z* (propagation step Δ*z* = *λ*_0_*/*4), and the computational window was padded to avoid circular-convolution artifacts. The complex fields *p*_*A*_(*x, y, z*) and *p*_*B*_(*x, y, z*) of the two sub-aperture transmissions were stored, and their coherent sum *p*_*A*_ + *p*_*B*_ was used to analyze the overlap geometry that drives the xAM response.

The three-event subtraction of Eq. 2 was not reproduced *in silico*. In experiment its diagnostic value comes from threshold-selective GV buckling where *p*_*A*_ +*p*_*B*_ *> p*_*b*_, whereas the solver was run as a linear, one-way forward propagator in which the subtraction is zero by construction. Nonlinear propagation is available in this solver [21] and in full-wave finite-difference methods [22], [23], but propagation nonlinearity alone does not produce the GV-specific residual. The simulations therefore use the linear pressure fields |*p*_*A*_ |,| *p*_*B*_ |, and |*p*_*A*_ + *p*_*B*_| to test whether a given (partition, *θ*) configuration satisfies the beam-confinement conditions on which the residual depends. The linear-field metrics defined below (peak intersection pressure, in-plane field uniformity, and side-lobe separation) serve only as *surrogate predictors* of nonlinear contrast, and their predictive value is tested *a posteriori* against the measured contrast-to-tissue ratio (CTR, defined in Sec. II-K; Sec. III-A, Fig. 4E).

**Fig. 4.**
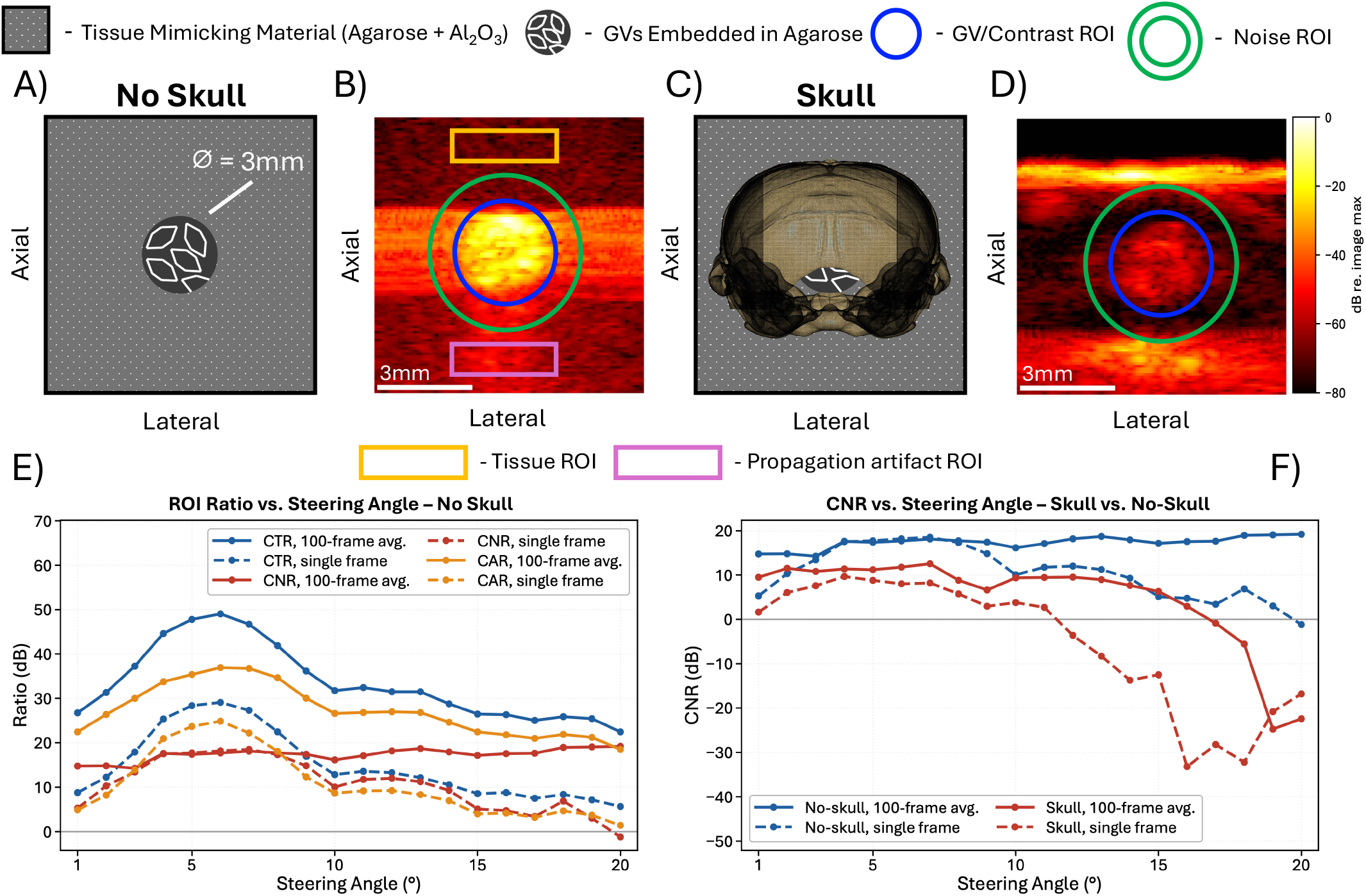
Phantom contrast quantification, no-skull versus through-skull. **(A)** No-skull tissue-mimicking phantom (agarose + Al_2_O_3_) with a 3 mm cylindrical GV inclusion. **(B)** Reconstructed xAM image of the no-skull phantom with the analysis regions of interest (ROIs): GV/contrast (blue), noise (green annulus), tissue (orange), and propagation-artifact (magenta). **(C)** Phantom with the *ex vivo* mouse skull placed in the beam path. **(D)** Reconstructed xAM image through the skull, with the gas-vesicle and noise ROIs only. The colorbar beside (D) applies to both the no-skull (B) and through-skull (D) images, in dB relative to the maximum of each image. **(E)** No-skull contrast-to-tissue (CTR), contrast-to-noise (CNR), and contrast-to-artifact (CAR) ROI ratios versus steering angle *θ*, for single frames and 100-frame averaging. **(F)** CNR versus steering angle *θ* for the skull and no-skull cases, for single frames and 100-frame averaging.

### E. Selection Criteria: Optimal Partition and Optimal xAM Angle

The ASM simulations were used to choose, *a priori*, both the partition and the steering angle *θ* for the experiments. Four metrics were defined: (1) the **peak-positive pressure** (PPP) at the intersection, 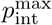, the maximum of |*p*_*A*_ + *p*_*B*_| on the bisector, which determines whether the GV buckling threshold is exceeded at the imaging depth; (2) the **X-wave beam width** *w(z)*, compared with the prediction *w* = *λ*_0_*/*(3 sin *θ*); (3) the **side-lobe separation** *d*_*SL*_, the lateral distance from the mainlobe peak to the nearest side-lobe peak in the bisector beam profile, compared with the prediction *d*_*SL*_ = *λ*_0_*/*(2 sin *θ*) of Favre and Maresca [3], which assumes monochromatic, far-field conditions; and (4) the **in-plane field uniformity**, the peak-to-trough ripple of the normalized pressure over the central *±* 3 mm of the in-plane profile (parallel to the bisector plane), a window twice the inclusion diameter, which measures how uniform the cross-propagation region is over the imaged target. Selection proceeded in two stages. The four partitions were first compared at a single steering angle on metrics (1) and (4), and the Y-split was chosen. The Y-split alone was then swept over *θ*, and metrics (1)–(3) were tracked against their predictions to fix the operating angle.

### F. Aberrating-Layer Simulations: One-Way Skull Propagation

The partition and angle studies above used a homogeneous medium. To model transcranial VxAM, the same solver was configured for heterogeneous propagation through a skull layer, with the model parameters listed in Table I. The skull geometry came from an openly distributed 4K micro-CT volume of a single adult *Mus musculus* dry skull (SCRI Bone and Skeletal Mechanics Laboratory, University of Washington; Bruker SkyScan 1272). This is a *different specimen* from the *ex vivo* skull used in the phantom measurements of Sec. II-H, and none of the acoustic properties in this subsection were measured on that physical specimen. Skull thickness, curvature, and suture geometry were taken directly from the scan, and the only geometric quantities we chose were those placing the transducer relative to the skull. The scan contains no soft tissue or brain and was acquired without a hydroxyapatite calibration phantom, so voxel intensities are raw, uncalibrated reconstruction counts.

The native stack was decimated eightfold to the *λ*_0_*/*4 solver grid by block-maximum downsampling, which preserves the thin cortical shell, rigidly aligned by principal-component analysis of the cortical-bone voxels, and cropped to the bone bounding box plus a 1 mm margin. Reconstruction counts *I* were mapped to sound speed and attenuation by a single piecewise-linear ramp between two histogram-derived intensity anchors (Table I), *t* = clip[(*I −* 5000)*/*20000, 0, 1], with *c* and *α*_0_ interpolated linearly in *t*. The upper sound-speed endpoint, *c*_*L*_ = 2425 m/s, is an effective value averaged over cortical and cancellous bone from broadband measurements on *ex vivo* murine calvaria [24] and was also used by Liang *et al*. to model the murine skull [25]. The upper attenuation endpoint, 8 dB cm^*−*1^ MHz^*−*1^, is the cortical-bone absorption of the International Transcranial Ultrasonic Stimulation Safety and Standards (ITRUSST) transcranial simulation benchmark (4 dB cm^*−*1^ at 500 kHz) [26], which we scaled linearly with frequency, and lies between the absorption and the total attenuation reported for skull bone [27]. Attenuation follows *α*(*f*) = *α*_0_*f* inside the skull and was not applied elsewhere, apart from the solver’s intrinsic *f* ^2^ water absorption, which is common to the homogeneous and heterogeneous conditions. The endocranial cavity, air-filled in the dry-skull scan, was segmented in each beam column as the non-bone region between the first and last bone voxel and assigned the soft-tissue properties of Table I. The resulting calvarial thickness, *≈* 0.3–0.4 mm, lies within the range reported for mice [24], [28]. The phase and amplitude screens carry no impedance term, so reflection loss is not represented and the simulated loss is a lower bound on the transmit-side loss through a real calvarium. The choice of intensity anchors, the attenuation endpoint, and their effect on the size of the simulated loss are detailed in the Supplementary Material.

The transmit field was normalized to a pre-skull peak-positive pressure of 250 kPa, so the simulation is interpreted in relative terms and its amplitude is not matched to the experimental *p*_int_ of Sec. II-C. No simulation in this work included nonlinear propagation; the medium was linear (*B/A* = 0, *β* = 1) in both the homogeneous and heterogeneous configurations. Heterogeneity entered the solver as phase and amplitude screens applied in the spatial domain between consecutive angular-spectrum steps, *φ* = 2*πf*_0_Δ*z* (1*/c−* 1*/c*_0_) and *A* = exp(*−α*_Np_Δ*z*) with *α*_Np_ = 11.51 *α*_0_*f*_MHz_ m^*−*1^ and *c*_0_ = 1540 m/s. The beam propagated dorsoventrally through the calvarium with about 1 cm of water between the transducer face and the dorsal bone surface, centered laterally on the endocranial cavity. The array was rotated 90° about its own axis so that its elevation (panel-stacking) direction ran anteroposteriorly, which places the two crossing plane waves in the anteroposterior–dorsoventral plane.

The main limitation of this approach, compared with a full heterogeneous-medium solver, is that it models only oneway forward propagation. It omits the second skull traversal of backscattered energy returning to the array, as well as internal reflections and multiple scattering in the bone microstructure. The one-way approximation is accurate for the transmit-side pressure distribution around the intersection but systematically underestimates degradation of the receive-side point-spread function. Simulated and experimental aberrated fields are therefore compared only in transmit quantities: peak intersection pressure, beam deflection, and beam broadening.

### G. Gas Vesicle Preparation and Phantom Construction

All phantom experiments used purified, stripped *Anabaena flos-aquae* (Ana) GVs, isolated from cultured *A. flos-aquae* by repeated hypertonic lysis and centrifugally assisted flotation as described previously [2], [29]. For nonlinear imaging, GvpC was removed by incubation in 6 M urea, which solubilizes the outer scaffolding protein and leaves the GvpA-based primary shell intact. Complete removal of GvpC was confirmed in a hydrostatic collapse chamber by monitoring optical density (OD) while the chamber was pressurized from 1 to 8 bar. The reported hydrostatic collapse midpoint of GvpC-stripped *Ana* GVs, 195 kPa applied pressure [29], lies well within this range. Stripped GVs have a lower collapse pressure and stronger second-harmonic scattering than GvpC-containing Ana GVs [29], [30]. GV concentration was measured by optical density at 500 nm (*OD*_500_) before phantom inclusion using a NanoDrop One spectrophotometer (Thermo Fisher Scientific, Waltham, MA, USA).

Phantoms were cast from agarose (2% w/v in degassed phosphate-buffered saline) with Al_2_O_3_ (0.2% w/v) for tissue-mimicking attenuation, in custom 3D-printed molds with a single 3 mm cylindrical GV inclusion **(Fig. 4A)**. A second design, used for the skull experiments, embedded an *ex vivo* skull (Sec. II-H) in the agarose before it set **(Fig. 4C)**.

### H. Ex Vivo Mouse Skull Specimen

The through-skull measurements used a single *ex vivo* skull from an adult male C57BL/6J mouse (approximately 6– 7 months old, body weight *≈* 33 g). All animal procedures were approved by the Institutional Animal Care and Use Committee of the University of North Carolina at Chapel Hill (protocol 25-145). The mouse was euthanized by CO_2_ overdose without transcardial perfusion. The calvarium was cleaned by gross mechanical dissection only, with no chemical or thermal maceration at any stage, so the bone was neither demineralized nor defatted. Skin and muscle were removed as completely as possible without compromising the skull. Removal of the periosteum could not be verified, because it was not resolvable by eye during dissection, and it is presumed to have been only partial. Likewise, some dura is presumed to have remained after the brain parenchyma was extracted. This residual periosteum and dura is an unquantified soft-tissue contribution to the through-skull acoustic path that the dry-skull simulation of Sec. II-F does not represent (Sec. IV-D). The skull was intact apart from a small posterior opening near the cerebellum, through which the brain parenchyma was removed. It was then stored fresh and unfixed in a sealed conical tube under refrigeration.

Before imaging, the specimen was submerged in water and held in a vacuum degassing chamber for 72 h to remove air trapped in the bone microstructure. The GV inclusion was then introduced into the cranial cavity through the posterior opening, at the approximate intracranial location of a glioblastoma tumor. The skull was embedded in agarose as in Sec. II-G (Fig. 4C), oriented so that the transmit beam crossed the intact skull before reaching the GVs. This specimen was used only for the phantom measurements and not for the simulation of Sec. II-F, whose skull properties were specified independently.

### I. Volumetric Acquisition: Electronic Steering vs. Robotic Translation

We compared two volumetric acquisition strategies, one in simulation and one in experiment. The *native* (electronic) strategy moves the xAM bisecting slab across the imaging volume by steering the two half-apertures to unequal angles (*θ*_*A*_ ≠ *θ*_*B*_), so that the plane waves cross at a lateral position set by the asymmetry Δ*θ* = *θ*_*A*_ + *θ*_*B*_. For a given two-half partition, the asymmetric sweep moves the intersection along the single axis normal to the partition line (Sec. III-E). The intended workflow used two passes from a fixed probe position, an asymmetric sweep with the X-split followed by one with the Y-split, so that together they would cover the lateral-elevation imaging plane without moving the probe.

The *native* strategy was simulated with ASM by sweeping Δ*θ* across the full *±* 4.8 mm extent of the probe. The bisector was considered coherent over the elevation range in which the element directivity still supported an efficient steered plane wave toward the target. Because the single-element directivity rolls off gradually, the edge of this range is also gradual (the resulting *≈* 50% elevation coverage is quantified in Sec. III-E and Sec. IV-E).

The *translated* (robotic) strategy **(Fig. 5)** fires the split sequence at a fixed elevation position (*Y* = 0) and builds the volume by moving the whole probe in elevation with an ABB IRB 1200 industrial robot arm (ABB Robotics, Zurich, Switzerland) driven by an IRC5 compact controller. A transistor–transistor logic (TTL) trigger pulse from the primary Vantage unit started each robot step once the frame batch at the current position had been acquired. The probe moved 200 *µ*m (*≈ λ*_0_) in elevation between acquisitions, with a 500 ms settling delay before each acquisition to let mechanical transients die out. A full volume was acquired at *≈* 1 s per position, excluding settling time and robot motion.

**Fig. 5.**
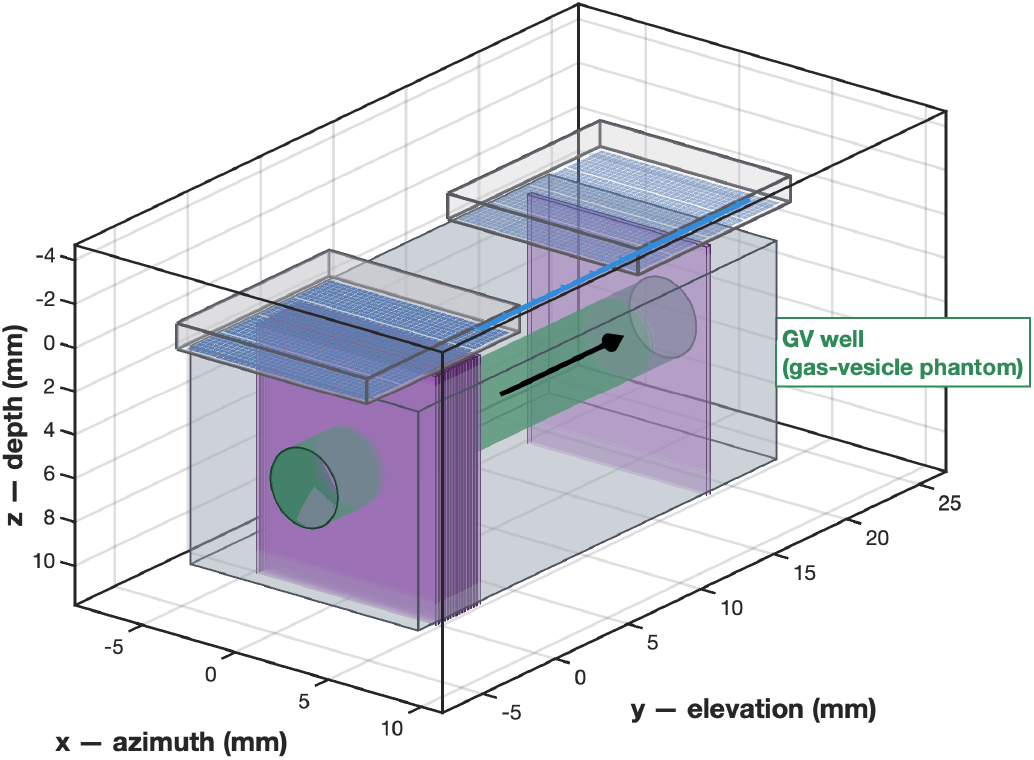
Robotic elevation-translation acquisition (*translated* strategy). A fixed-geometry xAM split sequence is fired at *Y* = 0 and the entire probe is translated in elevation by 200 *µ*m (*≈ λ*_0_) per step with an ABB IRB 1200 robot arm (IRC5 controller), triggered by the primary Vantage unit after each position’s frame batch. A typical sweep stacks 93 elevation positions spanning 18.1 mm into a coherent 3D VxAM volume.

### J. Image Reconstruction and xAM Signal Extraction

The xAM residual was formed on the raw per-channel echo data before beamforming. At each elevation position of the robotic sweep, the radio-frequency (RF) channel data from the three transmit events were combined according to Eq. 2. The residual was then beamformed onto a 3D slab with isotropic *λ*_0_*/*2 spacing by a custom in-phase/quadrature (IQ) delay-and-sum beamformer with geometric time-of-flight (geoToF) delays, which performs per-voxel delay-and-sum with integrated quadrature demodulation and returns a complex slab 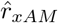 (**r**) directly, so no separate envelope detection was needed. The receive aperture for each voxel comprised the elements within a radius *z/*3 of it (an F-number of 3 applied to the radius), a diameter of 2*z/*3 and an effective F-number of 1.5. Each robot position was reconstructed over a 0.49 mm elevation extent (six planes at *λ*_0_*/*2 spacing), matched to the simulated bisector full width at half maximum (FWHM) at *θ* = 6° reported in Sec. III-D. The slab is narrower than the measured half-amplitude width of the transmitted sheet (616 *µ*m at 11.8 mm, Fig. S1), so that, with the 0.2 mm robot step, every voxel of the stitched volume is reconstructed only from positions whose bisector lies within*±* 0.25 mm of it, where the transmit field is within *≈* 4 dB of its peak. Slabs acquired at different probe positions share no common phase reference, so each was magnitude-detected before stitching. The stitch and the volume-domain frame average therefore operate on magnitude data, whereas the temporal averaging used for the contrast metrics (Sec. II-K) is coherent and precedes detection.

Because the robot moved the aperture 200 *µ*m between acquisitions (Sec. II-I), consecutive slabs overlap by *≈* 0.29 mm. To combine them into one volume, the slabs were placed at their robot positions on a common reconstruction grid, and each voxel of the final volume is the unweighted average of every slab that covers it (up to three).

### K. Image Quality Metrics

Each partition and angle was evaluated with three contrast ratios computed on the reconstructed xAM volume, all expressed in decibels: the *contrast-to-tissue ratio* (CTR), the *contrast-to-noise ratio* (CNR), and the *contrast-to-artifact ratio* (CAR). CTR and CAR are the ratios of the mean signal in the GV/contrast region of interest (ROI) to the mean signal in the tissue and propagation-artifact ROIs, respectively. CNR is the difference between the GV/contrast and noise ROI means, normalized by the standard deviation within the noise ROI:

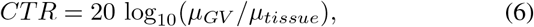

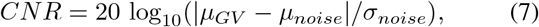

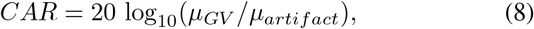

where *µ*_*GV*_ is the mean of 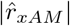 inside the GV/contrast ROI; *µ*_*tissue*_, *µ*_*noise*_, and *µ*_*artifact*_ are the means of 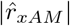 inside the tissue, noise, and propagation-artifact ROIs; and *σ*_*noise*_ is the standard deviation of 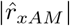 within the noise ROI, all defined on the reconstructed image **(Fig. 4B)**. The noise ROI is an annulus directly around the GV inclusion, so it contains the lateral point-spread-function side lobes of the X-wave bisector as well as uncorrelated noise. *σ*_*noise*_ is therefore not a pure noise estimate, and the reported CNR is conservative. Section III-B examines how this affects the response of CNR to temporal averaging. All three ratios were computed without the skull **(Fig. 4E)**. Through the skull only CNR was computed, so the comparison between the two cases uses CNR alone **(Fig. 4F)**. Every metric was evaluated for single frames and for 100-frame coherent temporal averages of the complex residual, formed before magnitude detection.

Volumetric resolution was measured with the circular-edge method [31], [32], applied to the cylindrical GV inclusion, whose axis lies along elevation so that its proximal surface, side walls and distal end cap present edges along the axial, lateral and elevation axes. The inclusion cross-section was located by a circle fitted to the 50% contour of the mean cross-section. Edge-spread functions (ESFs) were sampled along the boundary normal (axial and lateral, the latter within *±*20° sectors about horizontal to avoid the bright upper crescent and its acoustic shadow) or across the end cap (elevation), and averaged over the body of the inclusion. Each ESF was fitted with an error function in a fixed window anchored on its own 50% crossing [33], and the FWHM of the analytically differentiated fit, the line-spread function, is reported. Because the boundary has finite width and curvature, each FWHM is an upper bound. To test sensitivity to the boundary estimate, every fit was repeated with the circle center displaced by *±* 0.1 mm in lateral and depth.

## III. Results

All results come from a single proof-of-concept dataset: two tissue-mimicking phantoms (one plain and one containing the *ex vivo* skull), the scatterer-free ramp phantom of Sec. II-C, one GV preparation at *OD*_500_ = 3, and one *ex vivo* mouse skull. The candidate partitions were screened *in silico*, and the selected Y-split was then swept experimentally over steering angle. This scope was chosen to establish the feasibility of volumetric xAM on a matrix array. Generalization across GV concentrations, phantom geometries, and skull specimens is left for future work.

The transmit amplitude was calibrated before the partition and angle studies. Measured contrast rose steeply with drive amplitude, peaked at an intersection pressure of *≈* 600 kPa, and declined monotonically above it **(Fig. S2; Fig. S3 shows the full-aperture reconstruction discussed in Sec. IV-C)**. All subsequent no-skull acquisitions therefore used *p*_int_ *≈* 600 kPa, the amplitude of maximum contrast. The declining limb is not a collapse profile, and no collapse threshold is inferred from it. Section IV-C explains why and relates the operating point to previously reported collapse pressures, which the literature defines inconsistently.

### A. Partition Selection and Full Angle Sweep

The partitions were simulated in two stages: selection at a fixed reference angle (Fig. 3), then a full angle sweep on the selected partition (Fig. 6), whose predicted optimum was checked experimentally (Figs. 7 and 4E). In the first stage, the X-, Y-, and D-splits were compared at *θ* = 6° using their out-of-plane and in-plane profiles at a fixed depth **(Fig. 3C, D)**. Every split formed a coherent bisecting beam, but the Y-split gave the most uniform in-plane pressure. Its peak-to-trough ripple over the central *±* 3 mm was 1.0%, against 7.9% and 9.9% for the D-splits and 30.3% for the X-split **(Fig. 3D)**. Over the full bisection zone the D-splits are marginally flatter, because the Y-split has edge peaks beyond *±* 3.5 mm, but the *±* 3 mm window is the one matched to the inclusion imaged here. The Y-split also had the highest peak pressure 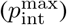, 1.00 at the bisector against *≈* 0.81 for the X-split and *≈* 0.79 for both D-splits **(Fig. 3C)**, and it was selected for the angle sweep.

**Fig. 6.**
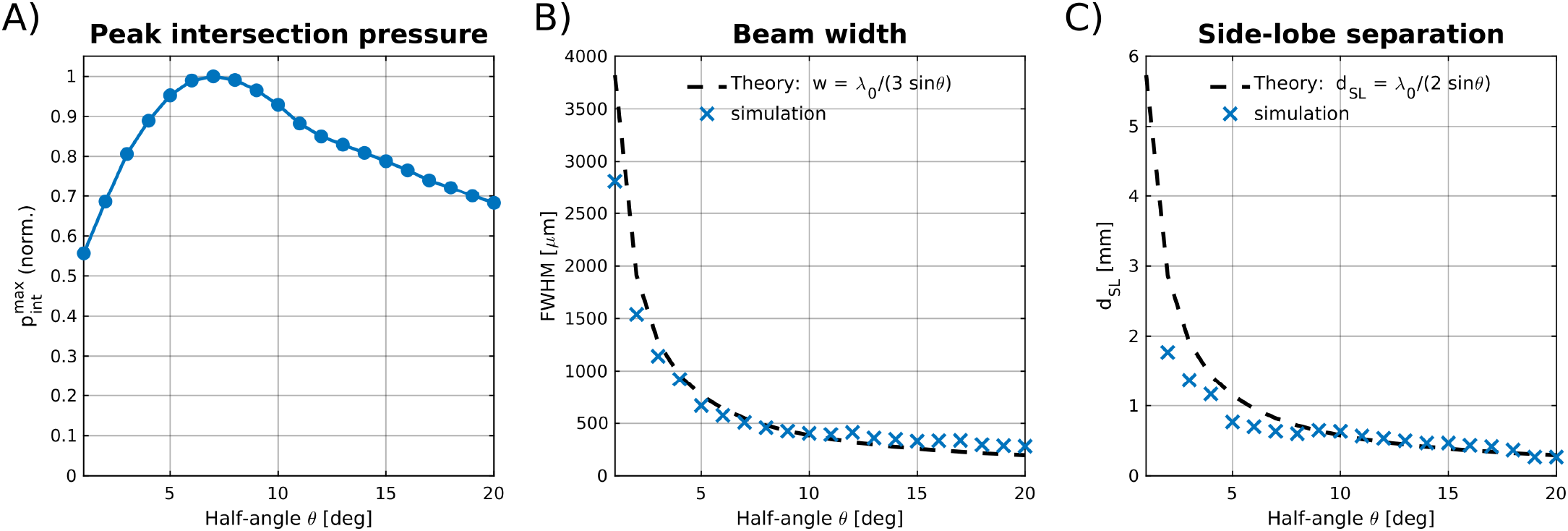
Y-split selection metrics versus steering angle *θ* (1°–20°) from homogeneous (no-skull) ASM simulation. **(A)** Normalized peak intersection pressure 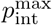. **(B)** Bisecting-beam width (FWHM): simulation (markers) versus the theoretical prediction *w* = *λ*_0_*/*(3 sin *θ*). **(C)** Main-lobe-to-side-lobe separation *d*_*SL*_: simulation (markers) versus the theoretical prediction *d*_*SL*_ = *λ*_0_*/*(2 sin *θ*).

**Fig. 7.**
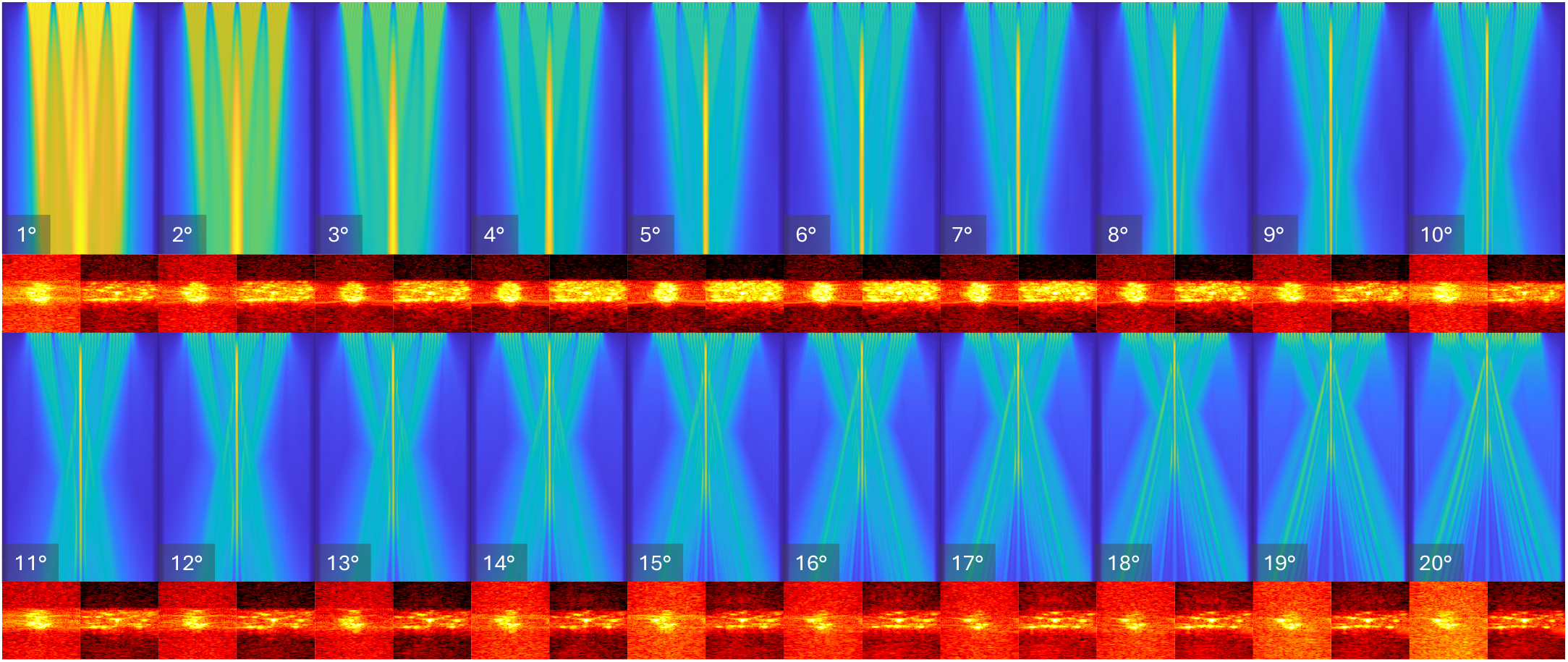
Steering-angle sweep on the Y-split partition over *θ* = 1°–20°: simulation versus experiment. For each angle, the simulated bisecting-beam pressure field is shown above the corresponding VxAM reconstruction of the gas-vesicle inclusion; angles 1°–10° are in the upper two rows and 11°–20° in the lower two rows.

In the second stage, the Y-split alone was swept over *θ ∈* [1°, 20°], and its out-of-plane and in-plane profiles were extracted at each angle. Peak intersection pressure, read from the out-of-plane profile at *z* = 11.8 mm, peaked near *θ* = 7°, with a nearly flat plateau from 6° to 8° (the differences among these angles were negligible) and a roll-off at larger angles as the beams’ overlap region shrank. The main-lobe-to-side-lobe separation *d*_*SL*_, computed from the same profiles, followed the theoretical scaling *d*_*SL*_ = *λ*_0_*/*(2 sin *θ*) and increased monotonically as *θ* decreased. It therefore has no interior optimum of its own. Instead it penalizes large steering angles, at which the side lobes close in on the main bisecting beam. The simulated beam width followed *w* = *λ*_0_*/*(3 sin *θ*) **(Fig. 6)**, but because it depends on *θ* alone it was the same for every partition and could not discriminate among them. At *θ* = 6° the side lobes were still well separated from the main beam and the peak pressure was on its plateau. Since the plateau is essentially flat from 6° to 8° and separability favors smaller angles, *θ* = 6° was adopted as the Y-split operating angle **(Fig. 6)**.

To check how well the simulations track the experimental beam physics, we acquired phantom xAM slabs at every angle in the range and compared the measured inclusion CTR and apparent main-beam width with the simulated predictions. The measured and simulated trends co-varied across *θ*, consistent with the selected operating angle of 6° **(Figs. 7 and 4E)**. The drive amplitude was held at the value calibrated at *θ* = 6°, however, so the *in situ* pressure at the inclusion follows 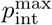 (*θ*) across the sweep, and this co-variation is only a consistency check. Unless noted otherwise, all later results use the Y-split at *θ*^*\**^ = 6°.

Propagation-artifact suppression was not one of the simulation metrics, because the linear formulation used here cannot produce the xAM residual (Sec. II-D) and so cannot predict the artifact. It was assessed experimentally instead. The contrast-to-artifact ratio measured over the same sweep **(Fig. 4E)** was *≈* 25 dB at *θ* = 6° for single frames. The artifact is thus suppressed but not eliminated at the selected angle, which lies well below the *θ >* 16.5° regime in which it is expected to nearly vanish [4]. The operating angle deliberately trades complete artifact suppression for the higher intersection pressure and larger side-lobe separation available at small *θ*.

### B. Contrast at a Fixed Probe Position

All contrast values in this subsection were measured at a single probe position, without robotic translation, during the steering-angle sweeps of Sec. III-A **(Fig. 4E, F)**. Without the skull **(Fig. 4E)**, the CTR at *θ* = 6° was *≈* 29 dB for single frames and *≈* 49 dB with 100-frame temporal averaging, and the CAR was *≈* 25 dB, rising to *≈* 37 dB. The frame-coherent component of the propagation-artifact ROI, estimated from the 100-frame average after removing the incoherent noise contribution, tracked the GV response across the steering-angle sweep and stayed 31–39 dB below it from 1° to 18°, above which it fell to the noise floor **(Fig. 4E)**. The artifact is therefore generated by propagation through the inclusion itself, and the decline of CAR at large *θ* reflects weaker GV excitation.

Averaging suppresses incoherent, frame-to-frame noise while the coherent GV residual persists, consistent with the incoherent nature of xAM image noise [3], which raised CTR by *≈*20 dB and CAR by *≈*12 dB. The tissue ROI mean fell by 20.0 dB under 100-frame coherent averaging, within 0.05 dB of the 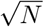 prediction for frame-uncorrelated noise. The tissue-mimicking material therefore contributes no detectable frame-coherent residual, which indicates that the three-event subtraction removes it to below the noise floor. One consequence is that the averaged CTR is set by the noise floor and is not directly comparable with *in vivo* values, whose denominator contains genuine tissue nonlinearity. The CNR, in contrast, was *≈* 18 dB with and without averaging. The tissue ROI that sets CTR is dominated by incoherent noise, but the CNR noise annulus is dominated by the coherent lateral point-spread-function side lobes (“whiskers”) of the X-wave bisector. These are spaced *d*_*SL*_ = *λ*_0_*/*(2 sin *θ*) *≈* 0.94 mm at *θ* = 6°, so they fall inside the annulus, and because they repeat from frame to frame, averaging does not reduce them.

The through-skull change in contrast is reported on CNR (Fig. 4F), which fell to *≈* 8 dB for single frames and recovered to*≈* 12 dB with 100-frame averaging. Through the skull, the coherent whisker structure fell below the random noise floor. Roughly half of the residual noise power was then incoherent and could be averaged down, which raised the through-skull CNR. Without the skull, no signal above the tissue ROI floor **(Fig. 4B)** appeared in the surrounding agarose, consistent with the subtraction working as intended on the unaberrated transmit/receive path. The through-skull acquisition was evaluated on CNR alone (Sec. II-K), so the same cannot be said for the aberrated path.

### C. Volumetric xAM Reconstruction

The Y-split at *θ* = 6° resolved the entire GV well (*OD*_500_ = 3) as a localized region of elevated xAM residual, with no visible elevation or lateral bleed into the surrounding agarose (Fig. 8A).

**Fig. 8.**
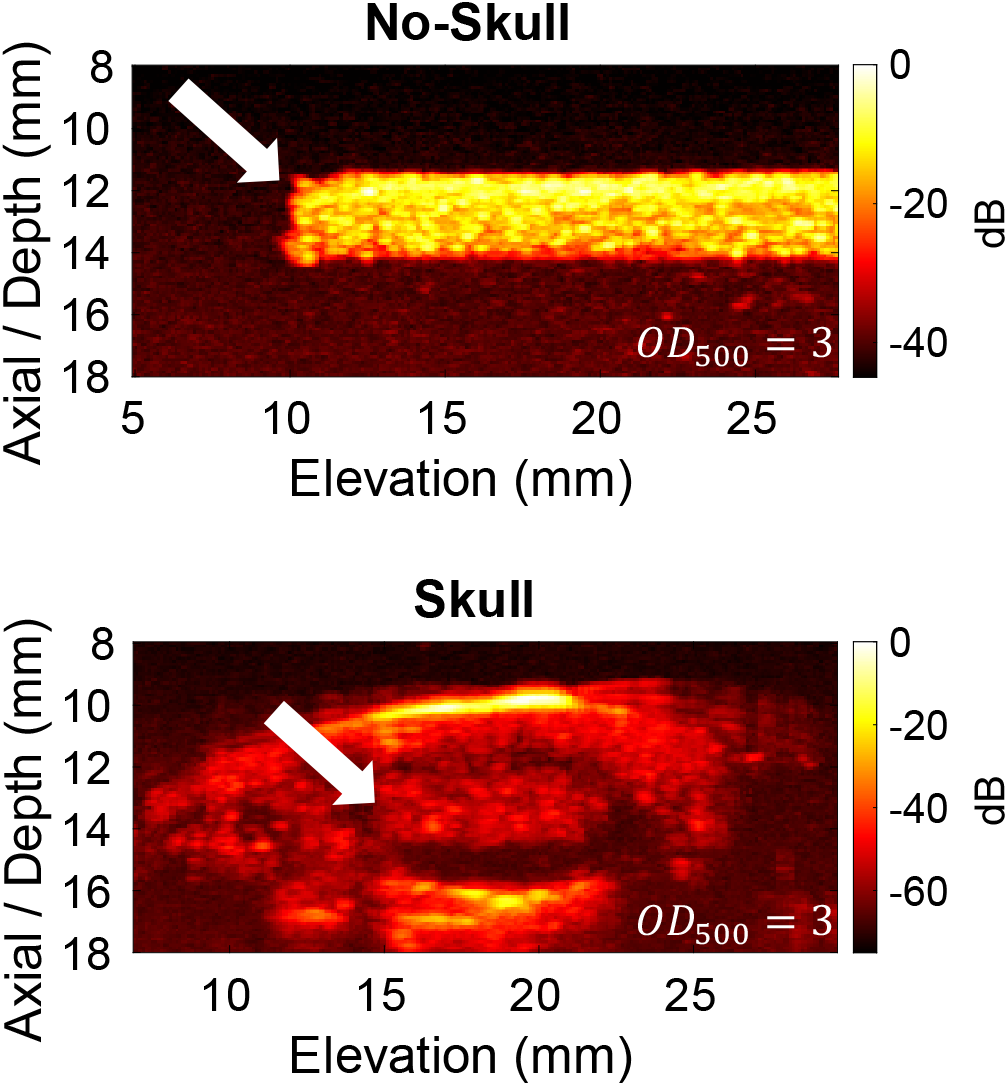
Volumetric VxAM reconstruction of the gas-vesicle inclusion (Y-split, *θ* = 6°). Stitched elevation-view maximum-intensity projection (MIP) of the reconstructed xAM residual (in dB) for the no-skull (top) and *ex vivo* through-skull (bottom) acquisitions at matched imaging depth (white arrows indicate the GV well; the bright superficial arc in the through-skull panel is the skull surface).

With the *ex vivo* mouse skull in the beam path and the GV well at the same depth, the GV signal was reduced but not eliminated. The well remained resolvable, with a broadened out-of-plane main lobe (Fig. 8B). We estimated the through-skull loss by cropping the reconstruction to a region matched to the resolved well in both conditions and comparing the mean xAM-residual amplitude (computed in the linear domain and expressed in dB) without and with the skull. The amplitude fell from 60.5 dB to 47.8 dB, a 76.8% reduction. Because the residual arises only where the bisector pressure exceeds *p*_*b*_, this loss is an indirect, GV-reported *estimate* of the skull’s effect on the *in situ* field that also includes return-path losses, and its mechanisms are separated in Sec. IV-D. This raw amplitude ratio is distinct from the contrast metrics of Sec. II-K.

### D. Simulation vs. Measured Skull-Induced Degradation

In the one-way ASM simulation of the Y-split at *θ* = 6° through the micro-CT mouse skull model, the peak intersection pressure fell by 1.27 dB **(Fig. 9)**, the bisector broadened from 0.49 to 0.59 mm FWHM (a factor of 1.20), and it was deflected by 0.10 mm in elevation. The off-bisector lobe at +3 mm, *≈* 0.53 of the main lobe without the skull, rose to *≈* 0.83 through it, while its *−*3 mm counterpart was essentially unchanged **(Fig. 9F)**. This simulated, forward-only pressure drop is far smaller than the measured 12.7 dB drop in residual amplitude (Sec. III-C), as expected: the simulation is one-way and linear, so it omits return-path attenuation, the shrinking of the above-threshold volume, and the skull’s own acoustic nonlinearity (Sec. IV-D, mechanism (d)), and the GV residual itself responds nonlinearly to incident pressure.

**Fig. 9.**
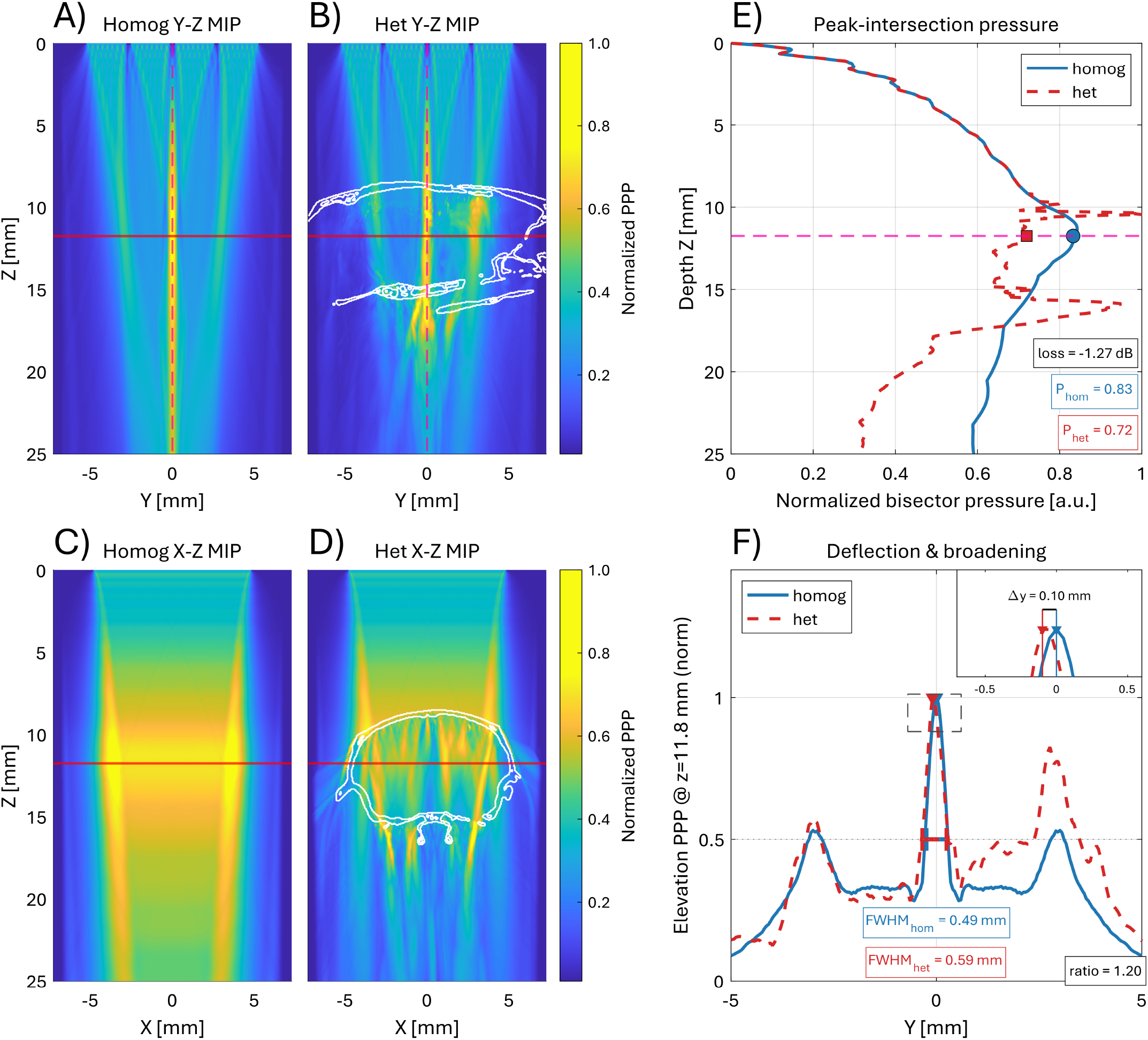
Simulated effect of the micro-CT mouse skull model on the VxAM bisecting beam (Y-split): homogeneous (no-skull) versus heterogeneous (through-skull) propagation. **(A, B)** Y–Z maximum-intensity projections (MIPs) of the bisector field in the homogeneous and heterogeneous media, respectively; the white contour outlines the skull. **(C, D)** Corresponding X–Z MIPs. **(E)** Peak-intersection bisector pressure versus depth for both media. **(F)** Elevation pressure profile at the analysis depth, showing the skull-induced beam deflection and broadening. Widths in (F) are measured with a non-interpolated half-level estimator, which is systematically*≈* 70 *µ*m narrower than the interpolated half-peak estimator used in Fig. 6(B). Absolute widths are therefore not directly comparable between the two figures, but the homogeneous-to-heterogeneous ratio reported in Sec. III-D is insensitive to the choice.

### E. Coherence Range of Electronic Asymmetric Angle Sweep

ASM simulations of the asymmetric sweep, with the two half-aperture angles set unequal (*θ*_*A*_ ≠ *θ*_*B*_), covered a *±*4.8 mm elevation range (*−*4.8 mm *≤ y ≤*+4.8 mm). The xAM bisector remained coherent only over the central half of the aperture, from *≈ −*2.4 mm to +2.4 mm (Fig. 10). Beyond this range the bisector region, and with it the xAM residual, broke down. A purely electronic asymmetric-angle sweep therefore covers only the central 50% of the swept elevation range coherently, and the outer 50% can be reached only by mechanical translation. The mechanism behind this limit and its consequences for volumetric acquisition are discussed in Sec. IV-E.

**Fig. 10.**
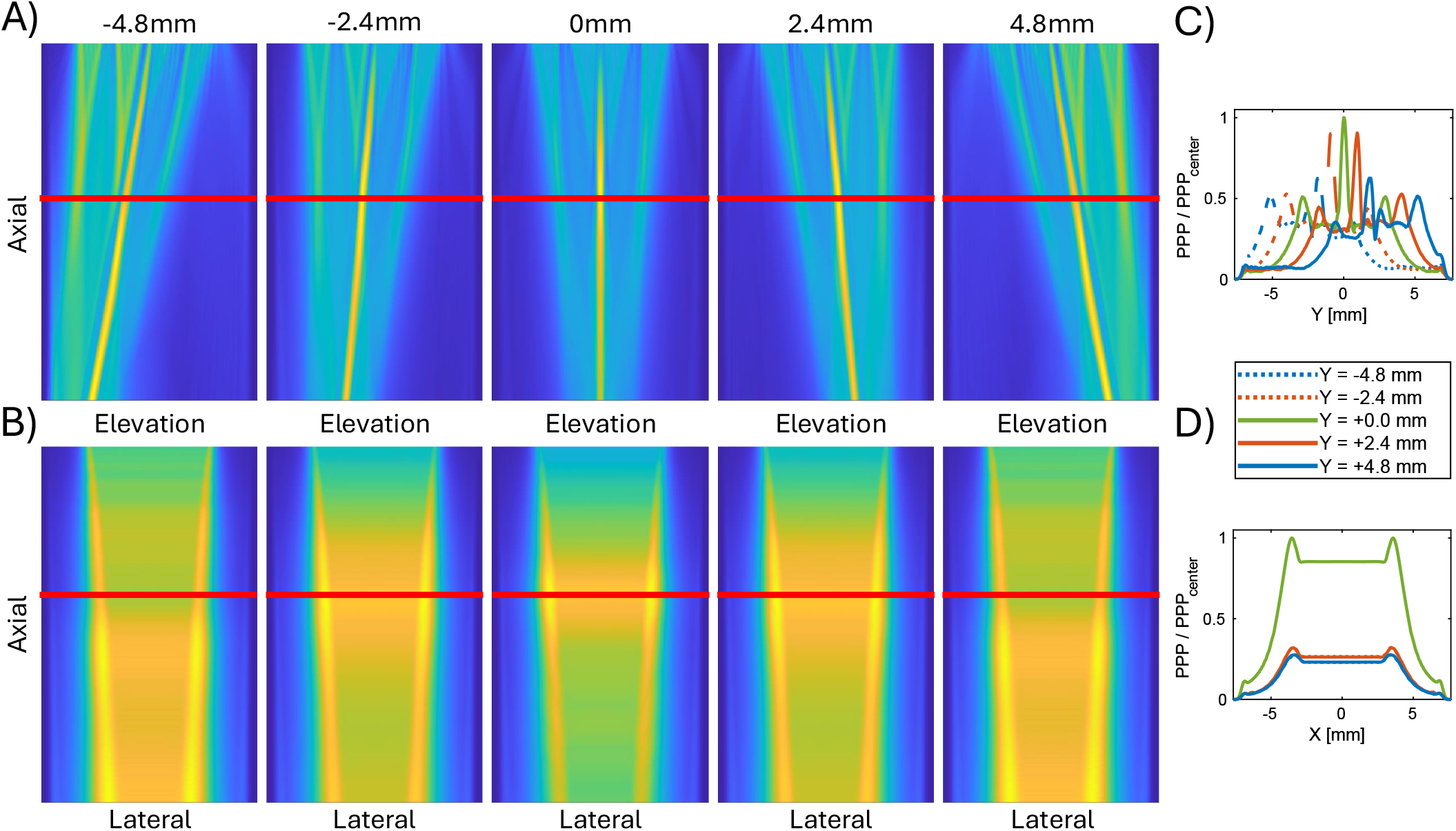
ASM-simulated electronic asymmetric-angle sweep across the elevation aperture, with the half-aperture angles made unequal (*θ*_*A*_ ≠ *θ*_*B*_) to steer the beam crossing to five elevation targets (*−*4.8, *−*2.4, 0, +2.4, +4.8 mm). **(A)** Axial–elevation bisector pressure fields at each steering target; the red line marks the 11.8 mm analysis depth. **(B)** Corresponding axial–lateral fields. **(C)** Normalized peak-positive-pressure (PPP) profiles along the elevation axis (*Y*) for each target. **(D)** Normalized PPP profiles along the lateral axis (*X*).

### F. Volumetric Resolution

Resolution was measured on the 200-frame-averaged no-skull volume by the edge-spread analysis of Sec. II-K **(Fig. S4)**. The fitted inclusion diameter was 2.63 mm, and ranges in parentheses give the spread over the *±* 0.1 mm center displacements; every value is an upper bound. The axial FWHM at the proximal surface is 348 *µ*m (347–354 *µ*m), compared with 281 *µ*m for half the spatial length of the measured emitted pulse. The emitted pulse is the appropriate reference because the transducer’s impulse response rings the one-cycle electrical drive out to about three cycles. The distal surface lies in the inclusion’s acoustic shadow and gives 705 *µ*m, a value set by attenuation through the inclusion and therefore excluded from the resolution estimates. The lateral FWHM is 400 *µ*m (399–509 *µ*m), with 346 and 468 *µ*m on the two walls, compared with *λ*_0_*z/D≈* 296 *µ*m for the receive aperture. The receive aperture (Sec. II-J) has diameter *D* = 2*z/*3 and lies inside the array along both *x* and *y* at the inclusion depth. This axis is set by receive beamforming and is not constrained by the crossing geometry, since *w* = *λ*_0_*/*(3 sin *θ*) governs only the out-of-plane axis. In elevation, which is both the out-of-plane crossing axis and the robotic stitching axis, the FWHM is 506 *µ*m (474–576 *µ*m). Elevation is coarser than lateral although the receive aperture is the same along both axes. The end-cap edge is measured across the stitched 0.49 mm slabs (Sec. II-J), and this acquisition does not separate the stitching from other contributions such as the shape of the cap. The contrast-to-noise ratio stayed above 6 dB wherever the inclusion was present, over at least 18.1 mm of elevation. Because the inclusion reached the edge of the reconstructed volume, these acquisitions give a lower bound on the usable elevation extent and do not characterize the field of view.

## IV. Discussion

### A. Interpretation of the Optimal Split

The main design rule from the partition analysis is *alignment with the non-emissive regions*. On a paneled matrix aperture, the best partition is the one whose bisector runs parallel to the inter-panel non-emissive regions. The Y-split satisfies this rule. Its bisector lies along the central non-emissive region between the four disjointed 32*×* 8 panels, over unpopulated material, whereas the X-split bisector runs perpendicular to these regions, across populated elements. Each X-split sub-aperture spans the full elevation and is interrupted by all three inter-panel non-emissive regions, while each Y-split sub-aperture spans two adjacent panels and contains a single internal non-emissive region. The more fragmented X-split aperture produces a less uniform field **(Fig. 3D)**. The Y-split is the orientation that Lee *et al*. excluded on a multiplexed version of the same array, expecting the gaps to disrupt the crossing waves [19]. On a fully addressed aperture, it instead gave the most uniform sheet and the highest peak pressure of all partitions.

Two further geometric effects support this alignment rule. The first is the sub-aperture baseline. The Y-split bisector lies parallel to the lateral *(x)* axis, so its two sub-apertures are separated along the longer, 10.6 mm elevation (*y*) dimension of the probe. Each Y-split sub-aperture is the upper or lower 16-row half of the 32*×* 32 grid, with a *y*-centroid at |*y* |*≈* 2.73 mm, giving a centroid separation of *≈* 5.46 mm along the bisector normal. The corresponding separation is 4.8 mm along 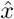 for the X-split and *≈* 4.83 mm for the D-splits. The crossing angle 2*θ*, the per-slice beam width *w* = *λ*_0_*/*(3 sin *θ*), and the side-lobe spacing are all set by *θ* alone, so every partition has the same beam cross-section at a given *θ*, and the partitions differ in baseline and in the orientation of their apertures relative to the non-emissive regions. Hydrophone measurements of the Y-split field confirm this scaling. At *θ* = 6° and the 11.8 mm analysis depth, the half-amplitude out-of-plane width is 616 *µ*m against 628.8 *µ*m predicted, and the side lobes fall at *±*950 *µ*m against the *±*943 *µ*m predicted by *d*_*SL*_ **(Fig. S1)**. The X- and D-split baselines are nearly equal (*≈* 4.80 and *≈* 4.83 mm, 0.7% apart, against 14% for the Y-split), so baseline distinguishes only the Y-split from the X/D pair. Any difference between the X- and D-splits must come from the orientation of their bisectors relative to the non-emissive regions and the grating-lobe axis.

The larger Y-split baseline lengthens the axial extent of the beam-overlap (X-wave intersection) region, in proportion to the baseline and inversely with tan *θ*, which sustains the crossing through the 11.8 mm analysis depth. The higher peak intersection pressure and more uniform sheet of the Y-split (Sec. III-A), however, come mainly from the *continuity* of its in-plane aperture. That aperture is in fact the shorter of the two (9.6 mm along *x* for the Y-split, against 10.6 mm along *y* for the X-split), but it is unbroken, whereas the in-plane aperture of the X-split is interrupted by all three inter-panel non-emissive regions. The benefit of alignment with the non-emissive regions is therefore aperture continuity, and it is large enough to outweigh a 9% shorter aperture.

The second effect is the periodicity the non-emissive regions impose on the aperture, with period 8*d* + *≈* 330 *µ*m = 2730 *µ*m 13.9*λ*_0_ along elevation only. Element-pitch grating lobes cannot distinguish the partitions. The 300 *µ*m lattice is the same in both axes, and the X- and Y-splits are related by a 90° rotation that leaves it unchanged. The panel periodicity breaks that symmetry. Because it runs along elevation, it lies *in* the X-split’s bisector plane but *normal to* the Y-split’s, and each X-split sub-aperture spans all four panels while each Y-split sub-aperture spans two. The X-split therefore carries a stronger periodic modulation in exactly the direction along which its sheet must be uniform, which is the far-field counterpart of the fragmentation described above. For the Y-split, the corresponding lobes arise from only two periods and fall *≈* 0.86 mm from the main lobe at the analysis depth, within 9% of the X-wave side-lobe spacing *d*_*SL*_ = *λ*_0_*/*(2 sin *θ*) = 0.94 mm, so they merge into the X-wave side-lobe structure.

### B. Contrast Advantage of Volumetric Matrix-Array xAM

The CTR values can be compared with those of 2D xAM on linear arrays [4] and of emerging volumetric nonlinear GV imaging methods such as sound-sheet microscopy [18] and Takoyaki ultrasound [19]. In a GV-loaded tissue-mimicking phantom at *OD*_500_ = 3, matrix-array volumetric xAM reached a well-to-tissue CTR of *≈* 29 dB for single frames and *≈* 49 dB with 100-frame averaging without the skull, and a CNR of *≈* 8–12 dB (single-frame to 100-frame-averaged) through the skull. For reference, Maresca *et al*. [4] reported an xAM CTR of *≈* 9 dB *in vivo* in the murine gastrointestinal tract (L22-14v at 15.6 MHz, *θ* = 19.5°). A direct numerical comparison is confounded by the *in vivo* versus phantom setting, the higher frequency, and the larger steering angle, so this value is cited only as a reference point. The architectural basis for a contrast advantage of the matrix array is discussed below.

This implementation complements the other volumetric non-linear GV imaging methods (Table II). NSSM covers a volume by sweeping a thin acoustic sheet through it and obtains contrast from an xAM residual. Takoyaki ultrasound acquires volumetric data by scanning multiple discrete focal points with a multiplexed matrix array. Compared with harmonic filtering, xAM has two qualitative advantages. First, the three-event subtraction removes linear arrivals from the image regardless of their spectral content, whereas harmonic filtering relies on clean spectral separation between the fundamental and the second harmonic [14], which becomes less reliable at higher pressures, where harmonic broadening of the fundamental spills into the passband and erodes the contrast signal. Second, provided the per-element drive is identical across the three events, transmit-side nonlinearities such as amplifier saturation enter all three events equally and should cancel in the sub-traction, whereas they add harmonic content that a harmonic filter cannot separate from GV signal. The cost is that each xAM frame needs three coordinated transmit-receive events (two half-aperture and one full-aperture) instead of the one or two used in harmonic or AM imaging.

**TABLE II.**
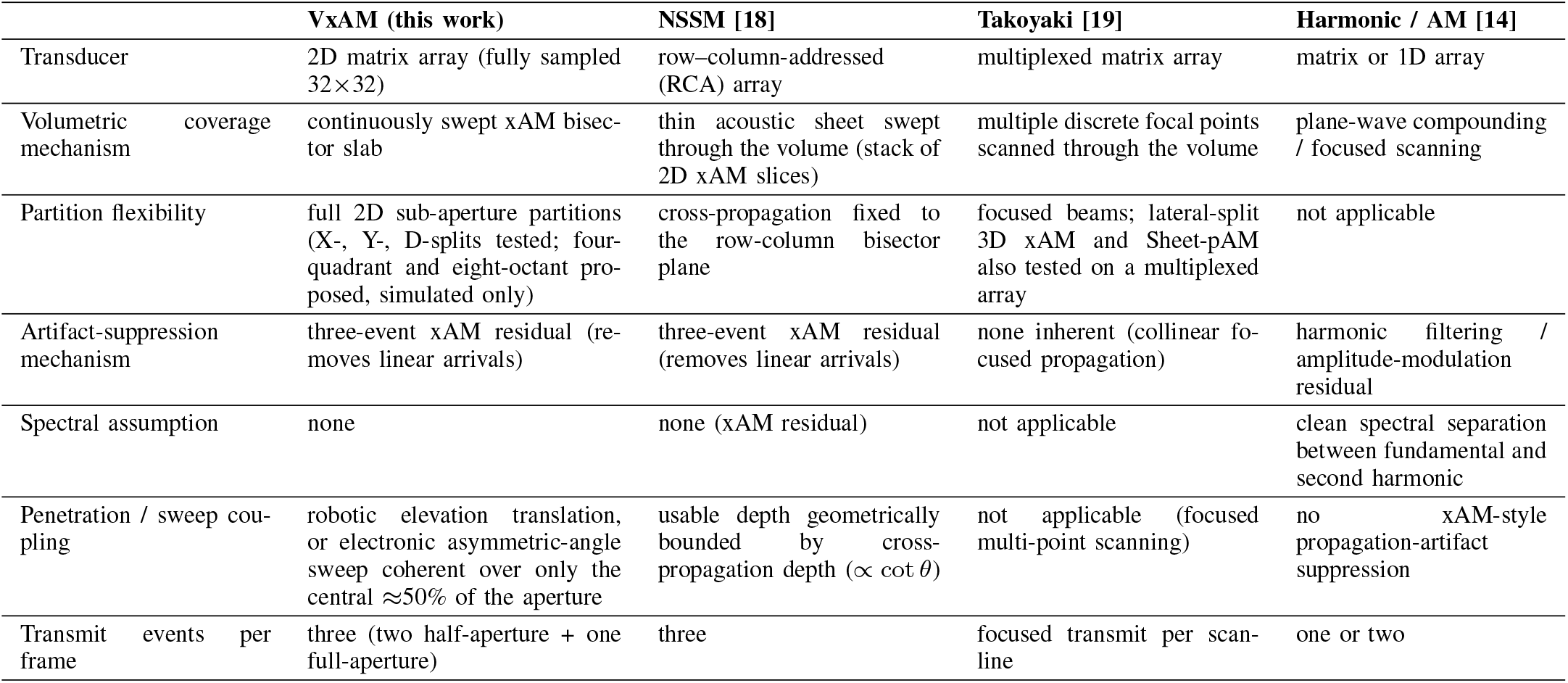
Comparison of Volumetric Nonlinear Gas-Vesicle Imaging Approaches.

|  | **VxAM (this work)** | **NSSM [18]** | **Takoyaki [19]** | **Harmonic / AM [14]** |
|---|---|---|---|---|
| Transducer | 2D matrix array (fully sampled 32×32) | row–column-addressed (RCA) array | multiplexed matrix array | matrix or 1D array |
| Volumetric coverage mechanism | continuously swept xAM bisec-tor slab | thin acoustic sheet swept through the volume (stack of 2D xAM slices) | multiple discrete focal points scanned through the volume | plane-wave compounding / focused scanning |
| Partition flexibility | full 2D sub-aperture partitions (X-, Y-, D-splits tested; four-quadrant and eight-octant pro-posed, simulated only) | cross-propagation fixed to the row-column bisector plane | focused beams; lateral-split 3D xAM and Sheet-pAM also tested on a multiplexed array | not applicable |
| Artifact-suppression mechanism | three-event xAM residual (re-moves linear arrivals) | three-event xAM residual (removes linear arrivals) | none inherent (collinear fo-cused propagation) | harmonic filtering / amplitude-modulation residual |
| Spectral assumption | none | none (xAM residual) | not applicable | clean spectral separation between fundamental and second harmonic |
| Penetration / sweep cou-pling | robotic elevation translation, or electronic asymmetric-angle sweep coherent over only the central ≈50% of the aperture | usable depth geometrically bounded by cross-propagation depth (∝ cot θ) | not applicable (focused multi-point scanning) | no xAM-style propagation-artifact suppression |
| Transmit events per frame | three (two half-aperture + one full-aperture) | three | focused transmit per scan-line | one or two |

This approach differs from prior nonlinear GV imaging in how its interaction region is formed. Conventional xAM on a 1D linear array confines the cross-propagating plane waves to the imaging plane, with the elevation dimension fixed by an acoustic lens. The RCA array used in NSSM forms its cross-propagation from orthogonal row and column groups that meet along the row–column bisector. In both cases, the nonlinear interaction region is a thin *sheet* that is swept through the volume, mechanically for a linear array and electronically for an RCA array. On a fully sampled 2D matrix array, lateral and elevation elements are independently addressable, so a half-aperture split aligned with the non-emissive regions launches two large 2D plane waves that intersect across the full lateral extent of the array in a single transmission. The resulting bisector is also thin in elevation (*≈* 0.5 mm; Sec. III-D) and broad laterally, which we refer to as a slab. What differs is how it is formed: *coherently*, in a single transmission, by two independently steerable 2D half-apertures that are free of the row–column constraints of Sec. I and can be split along any of the partition directions of Sec. II-B.

Robotic translation is also the usual way to build volumes with a 1D linear array, by stepping the probe in elevation and stacking 2D images, but the two arrays sample elevation differently. A 1D array has no electronic control in elevation. Its slice thickness is set by a fixed acoustic lens, is narrowest only near the lens focus, and each 2D image integrates the echo signal across that thickness. A stack of such images therefore has an elevation resolution equal to the local slice thickness, which is coarser than the in-plane resolution and changes with depth, and an elevation sampling equal to the translation step. Its voxels are correspondingly anisotropic, fine in the image plane and equal to the translation step in elevation. On the matrix array, elevation is beamformed. Each robot position yields a slab resolved into *λ*_0_*/*2 planes (six planes over 0.49 mm, Sec. II-J) with receive focusing along *y* as well as *x*, and the 0.2 mm step overlaps consecutive slabs so that each voxel is averaged over up to three positions. The stitched volume is sampled on an isotropic *λ*_0_*/*2*≈* 99 *µ*m grid in all three axes, finer than the measured resolution along each of them, so the voxel size does not limit resolution in any direction. Translation then extends coverage, while elevation resolution comes from the beamformer and the crossing geometry. The measured elevation FWHM, 506 *µ*m, is 26% coarser than the lateral 400 *µ*m (Sec. III-F). For xAM the two cases also differ in what confines the nonlinear signal along the translation axis. On a linear array the cross-propagation plane is the imaging plane, so only the lens limits the residual in elevation. For the Y-split, the crossing itself confines the interaction to the *≈* 0.5 mm bisector along the translation axis, and receive beamforming resolves it within that slab.

For contrast, the X-wave beam cross-section depends only on the steering angle (*w* = *λ*_0_*/*(3 sin *θ*)) and is the same for every partition (Sec. IV-A), so any benefit of the matrix-array geometry comes through the peak intersection pressure and field uniformity at a fixed cross-section. Here those were provided by the unbroken in-plane aperture, aligned with the non-emissive regions, and the larger sub-aperture baseline of the Y-split. Tighter focusing plays no part, since true 3D concentration occurs only for the higher-order Bessel-like decompositions of Sec. IV-F. Because GV buckling is threshold-gated and supralinear in incident pressure, a higher and more uniform *in situ* pressure both enlarges the buckling-active volume (more of the intersection exceeds *p*_*b*_) and raises the per-voxel residual amplitude where *p > p*_*b*_, and each of these increases the residual that defines the contrast.

The higher and more uniform intersection pressure should therefore favor GV detection, but the phantom CTR reported here (*≈* 29 dB single-frame, *≈* 49 dB averaged) does not isolate that effect. The measured value is dominated by experimental factors: a clean phantom, the high *OD*_500_ = 3 concentration, low-threshold stripped *Ana* GVs, and the *≈* 20 dB gain from 100-frame coherent averaging (Sec. III-B). The aperture geometry of the fully sampled matrix array is therefore a qualitative advantage that may contribute to the elevated contrast but does not account for it.

Temporal averaging raised the no-skull CTR and CAR by*≈* 20 and *≈* 12 dB without changing the three-event transmit sequence, but only to the extent that the noise is incoherent (Sec. III-B). In the transcranial setting it therefore complements the aberration robustness of the three-event subtraction without replacing it (Sec. IV-D).

### C. Operating Pressure and the Definition of Collapse

The peak-positive pressure used throughout this work (*p*_int_ *≈* 600 kPa, Sec. II-C) is close to values that have been reported as the acoustic collapse threshold of stripped *Ana* GVs, so we compare them explicitly here. “Collapse pressure” is a percentile of a distribution, and the literature does not choose that percentile consistently. Collapse is usually quantified by fitting the normalized pressure-sensitive optical density (hydrostatic) or backscattered amplitude (acoustic) to a Boltzmann sigmoid, 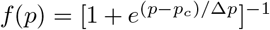, where *f* is the intact fraction of the population, *p*_*c*_ the collapse midpoint, and Δ*p* the width [5], [6], [29]. Reported values are almost always the midpoint *p*_*c*_, but first-disappearance and cumulative pulse-train criteria are also used [5], and these criteria are not interconvertible.

The choice of percentile moves the quoted threshold sub-stantially. For the stripped *Ana* preparation used here, Lakshmanan *et al*. report an acoustic collapse profile at 6.25 MHz with *p*_*c*_ = 571 kPa and Δ*p* = 14.5 kPa [29]. Propagating these parameters through the sigmoid, the same preparation and dataset give a nominal collapse pressure of *≈* 539 kPa if collapse is declared at 10% of the population, 571 kPa at the 50% midpoint, *≈* 614 kPa at 95%, and *≈* 638 kPa at 99%, a spread of *≈* 100 kPa due entirely to the definition. We adopt no collapse criterion here, since the operating amplitude was chosen by maximizing measured contrast (Sec. II-C). The operating point of *≈* 591 kPa lies between the 50% and 95% points of this profile, and the calibration step is comparable to the spacing between those percentiles.

The measurement geometry gives a further reason not to read the pressure ramp of Sec. III as a collapse curve: two thresholds operate at once across the inclusion. The xAM residual arises only where the bisector pressure exceeds the buckling threshold *p*_*b*_, since reversible shell buckling is what produces the nonlinear scattering (Sec. I). Irreversible collapse, which removes a GV from the population, requires a still higher pressure. Neither contour is fixed: raising the drive widens the supra-threshold cross-section of the main bisecting lobe and lifts the X-wave side lobes, spaced *d*_*SL*_ = *λ*_0_*/*(2 sin *θ*) *≈* 0.94 mm at *θ* = 6°, above *p*_*b*_ in their own right, so both contours expand together. The collapse contour advances through the core of the inclusion, removing GVs that had been contributing signal, while the buckling contour advances into the periphery, recruiting GVs that had been below *p*_*b*_ and so had neither contributed signal nor been exposed to a destructive amplitude. The measured curve is therefore the convolution of these two processes. This explains both its shape, a gradual monotonic decline where a uniformly exposed population would give a sigmoid, and why complete signal loss is deferred to pressures well above the maximum: the collapse contour must sweep past the full extent of the inclusion before every GV has been removed. Our data show this effect directly. Reconstructing the same voltage sweep over the full aperture instead of the bisector slab increases the elevation extent of the analysis volume from 0.49 to 9.96 mm, admitting roughly five orders of X-wave side lobes on each side where the slab admits none. The apparent optimum moves from*≈* 600 kPa in the bisector-restricted reconstruction to *≈* 1469 kPa in the full volume **(Figs. S2 and S3)**, because off-bisector GVs brought above *p*_*b*_ by the side lobes add to the contrast metrics without contributing to the intended bisecting slab. The bisector-restricted value is the relevant one and is the basis for the operating point used here.

Two further differences separate our operating condition from published values. Acoustic and hydrostatic collapse pressures are not interchangeable, the acoustic collapse pressure exceeding the hydrostatic midpoint by *≈*2.9*×* for stripped *Ana* [29] and by *≈* 9*×* for *Halo* GVs [5], so the hydrostatic test of Sec. II-G is a quality check on GvpC removal and not a bound on the acoustic operating point. In addition, no collapse measurement has been reported at 7.81 MHz, although the *Halo* threshold shows no decrease toward lower frequencies between 27.5 and 12.5 MHz [5] and the nonlinear GV response is expected to vary little above *≈* 1 MHz, where little gas crosses the shell during compression [30].

All metrics reported here are steady-state values. At each amplitude the inclusion was insonified until the residual stabilized before the averaging batch was acquired (Sec. II-C), and over the 100-frame average the mean residual in the GV ROI changed by 0.09 dB, so repeated imaging at a fixed amplitude is non-destructive in the steady state, which is the condition averaging requires. The protocol does not show whether GVs collapsed during the preceding stabilization interval (Sec. IV-G). Two published observations are consistent with stable imaging near the collapse midpoint. In ARG-expressing bacteria the xAM signal decays above *≈* 1 MPa to a nonzero plateau, still holding 57% of its initial amplitude at 2.23 MPa [8], and the original xAM implementation used single-cycle transmits on urea-stripped *Ana* GVs without reporting collapse [4]. The collapse thresholds cited here, however, were measured with four-cycle [29] and six-cycle [5] bursts. Linear (eigenvalue) and explicit multi-cycle dynamic buckling analyses of an *Ana* GV agree to within 1 kPa (332 versus 331 kPa) [34], so we do not expect the shorter burst to raise the threshold, but our transmit condition lies outside the range over which those thresholds were established. Measuring the full collapse profile of this preparation at 7.81 MHz would remove the remaining extrapolation.

## D. Implications for Transcranial Imaging

With the *ex vivo* skull in place, the mean xAM-residual amplitude fell by 12.7 dB, but the GV well remained resolvable (Sec. III-C). Both observations follow from how the three-event subtraction works. The linear through-skull propagation terms enter all three transmit events identically and cancel in the subtraction whether or not the incident wavefronts were aberrated on the way, so the aberrated linear background is removed and the contrast mechanism itself survives a stationary aberrating layer. Because the well remained resolvable through bone, with tissue suppressed on the unaberrated path, this result is a step toward transcranial molecular imaging in rodents. The subtraction cannot, however, preserve amplitude. The residual arises only where the bisector pressure exceeds the buckling threshold *p*_*b*_, so its magnitude is set by the *in situ* field, and anything that lowers the field at the inclusion lowers the residual with it. The measured 12.7 dB loss therefore reflects how much the skull reduced the residual, mainly by reducing the *in situ* field, and is not evidence that the subtraction failed. The distinction matters for translation, because the two would call for different remedies.

Four mechanisms contribute to the loss, and the subtraction removes none of them. The first three are expected to dominate. (a) After one-way skull traversal, the incident pressure is reduced by insertion loss and diffuse scattering in the cranial bone. Part of the designed intersection volume then falls below *p*_*b*_ and generates no xAM residual even if the subtraction is otherwise perfect. This predicts that the buckling-active volume shrinks while the per-voxel xAM contrast is preserved in regions still above *p*_*b*_. (b) The aberrated wavefront from each half-aperture is no longer a clean plane wave at the intended steering angle *θ*. The two beams therefore cross in a slightly displaced and distorted intersection region with less main-lobe overlap, and some of the transmit energy is redirected into grating lobes and side lobes outside the intended bisecting volume. This predicts a broadening and elevation shift of the xAM well, qualitatively consistent with the results above. (c) On the return path, the skull attenuates the higher-frequency content of the buckling-generated nonlinear signal more than the fundamental, because attenuation in cranial bone increases with frequency (Sec. II-F). Less of the buckling-generated energy therefore reaches the array than the one-way forward simulation would suggest. This predicts that the xAM residual scales uniformly with its spatial distribution preserved. (d) Cranial bone, like any real solid, is also acoustically nonlinear. As with the cumulative propagation nonlinearity of Sec. I, the contribution generated along each half-aperture path separately cancels in the subtraction. What survives is the cross-term*∝ p*_*A*_*p*_*B*_ generated where the two beams overlap inside the bone [12], [35], which adds a GV-independent background to the residual without scaling or displacing the GV signal. This term should be small, since nonlinear distortion accumulates with propagation distance and the calvarium modeled here is only*≈* 0.3–0.4 mm thick, against a centimeter-scale water and gel path. We include mechanism (d) for completeness; it is not expected to be a leading contributor, and it is absent from the simulation, which was linear (Sec. II-F).

Mechanisms (a)–(d) are all properties of the bone, but the specimen retained an unquantified amount of periosteum and dura (Sec. II-H), and the aberrating-layer model was built from a dry-skull micro-CT volume with no soft tissue at all (Sec. II-F). Residual soft tissue would add attenuation and aberration on both the transmit and return paths, contributing to (a) and (c) without being represented in the simulation, and is there-fore a candidate contributor to the gap between the simulated and measured losses of Sec. III-D. The relative weight of (a)–(d) can be tested in future work by repeating the measurement at several transmit pressures (isolating the buckling-threshold mechanism), comparing simulated and measured side-lobe energy (isolating beam redistribution), comparing the measured and simulated spectra of the subtracted residual (isolating return-path attenuation), and acquiring the three-event sequence in a GV-free skull preparation (isolating the skull self-nonlinearity background).

### E. Electronic Asymmetric-Angle Sweep

As described in Sec. II-I, the *native* electronic sweep moves the xAM bisector region in elevation without moving the probe, by steering the two half-apertures to unequal angles (*θ*_*A*_ ≠ *θ*_*B*_). On the fixed matrix aperture this relative angle is the only electronic steering mechanism available, unlike the equal-angle sweep used on 1D linear arrays, which walks the sub-aperture across the array [4]. The limited coherent range reported in Sec. III-E, the central half of the aperture (Fig. 10), is explained by the finite steerability of the individual Vermon elements. The element width *w*_e_ = 280 *µ*m is only *≈* 1.4*λ*_0_ at *f*_0_ = 7.81 MHz, which gives a narrow single-element directivity with a *−*3 dB half-width of *≈*18°. Measured from the array center to the target at the analysis depth, reaching the swept edge (*y*_*c*_ = *±*4.8 mm) requires a steering angle of *≈*22°, against *≈*11° for *y*_*c*_ = *±* 2.4 mm. The corresponding single-element responses are *≈−* 4.6 and *≈−* 1.2 dB, so the edge targets lie beyond the *−*3 dB half-width and the mid-aperture targets within it. Because the roll-off is gradual (Sec. II-I), the taper lowers transmit efficiency progressively toward the aperture edges. The angles required of the individual half-apertures are larger still, since each sub-aperture is offset in elevation from the array center. As quantified in Sec. III-E, each asymmetric-angle sweep therefore recovers only the central half of the swept elevation range, and the rest can be reached only by mechanical translation. Covering the full lateral-elevation plane without moving the probe would require pairing an X-split asymmetric-angle sweep (steering across *x*) with a Y-split sweep (steering across *y*). Section III-A, however, identified a single best partition, and this two-pass workflow would give half of its volumetric coverage to the X-split, with poorer field uniformity in elevation. For these reasons, all experimental volumes were acquired by robotic probe translation (Sec. II-I). Stepping a cylindrically focused sheet electronically along elevation, as in Sheet-pAM [19], avoids this steering limit, but its collinear AM transmissions do not provide the propagation-artifact suppression of xAM.

The coherent range established here for the Y-split also limits how much of the volume any purely electronic 3D xAM mode could cover without mechanical help on an aperture the size of this Vermon probe. Hardware routes to a wider range include finer-pitch matrix arrays, whose smaller elements have broader directivity, and hybrid acquisitions that combine a coarse mechanical sweep with fine electronic asymmetry to recover acquisition speed without losing coverage. These trade-offs matter most for *in vivo* use. The robotic workflow used here is slower than a full-aperture electronic mode would be, and therefore more exposed to physiological motion, cardiorespiratory gating artifacts, and inter-slab registration error, whereas a purely electronic mode would need no probe motion at all. For rodent brain imaging, the volume acquisition time reported here, on the order of one to two minutes (Table I), would require the animal to be anesthetized with its head fixed in a stereotaxic frame.

### F. *Higher-Order Decompositions, Monolithic Arrays, and* In Vivo *Outlook*

This work was restricted to the three primary sub-aperture partitions (X-, Y-, and D-splits). The matrix aperture also supports two higher-order decompositions that neither a 1D linear array nor a row–column-addressed array can form: a four-quadrant partition combining the X- and Y-splits, fired as two opposing quadrant pairs, and an eight-octant decomposition. In preliminary ASM simulations, both produce Bessel-like conical beams in the central intersection region, and the octant decomposition reaches a higher peak pressure than the four-quadrant one. We leave their full characterization to future work.

Testing these configurations experimentally would not change the number of transmit events per frame. The three-event subtraction (Eq. 2) is unchanged, and the higher-order configurations are defined entirely by the per-element delay profiles of each sub-aperture. For the four-quadrant partition, the X-split and Y-split delay profiles are superimposed so that the sub-aperture A transmit drives one pair of opposing quadrants and the sub-aperture B transmit drives the other. A single pair of half-aperture transmits then imposes both bisections at once and forms a Bessel-like beam. The octant partition is built the same way from the X-, Y-, and D-split profiles, with the two half-aperture transmits driving complementary groups of four octants. In both cases, the per-frame acquisition time, the amount of data entering the subtraction, and the inter-scanner synchronization requirements are the same as for the baseline xAM sequence.

A further hardware option is the recently released monolithic Vermon matrix array, which has no inter-panel non-emissive regions. The Y-split’s advantage over the X-split on the paneled probe comes from two coupled asymmetries identified in Sec. IV-A: the alignment of its bisector with the non-emissive regions and its larger sub-aperture baseline (*≈* 5.46 mm versus *≈*4.8 mm). Both are specific to the paneled aperture. On a monolithic *square* aperture, removing the non-emissive regions eliminates the asymmetry they introduce and equalizes the X- and Y-split baselines at the same time. The *x/y* symmetry is restored, and the X- and Y-splits become equivalent by construction (on a non-square monolith the baselines would still differ). The robotic-translation workflow used here then extends directly to a two-direction sweep, with the X-split translated along one axis and the Y-split along the orthogonal axis, covering the full lateral-elevation plane without rotating the probe between passes and without the penalty that the paneled aperture imposes on any X-directed sweep **(Fig. 11)**.

**Fig. 11.**
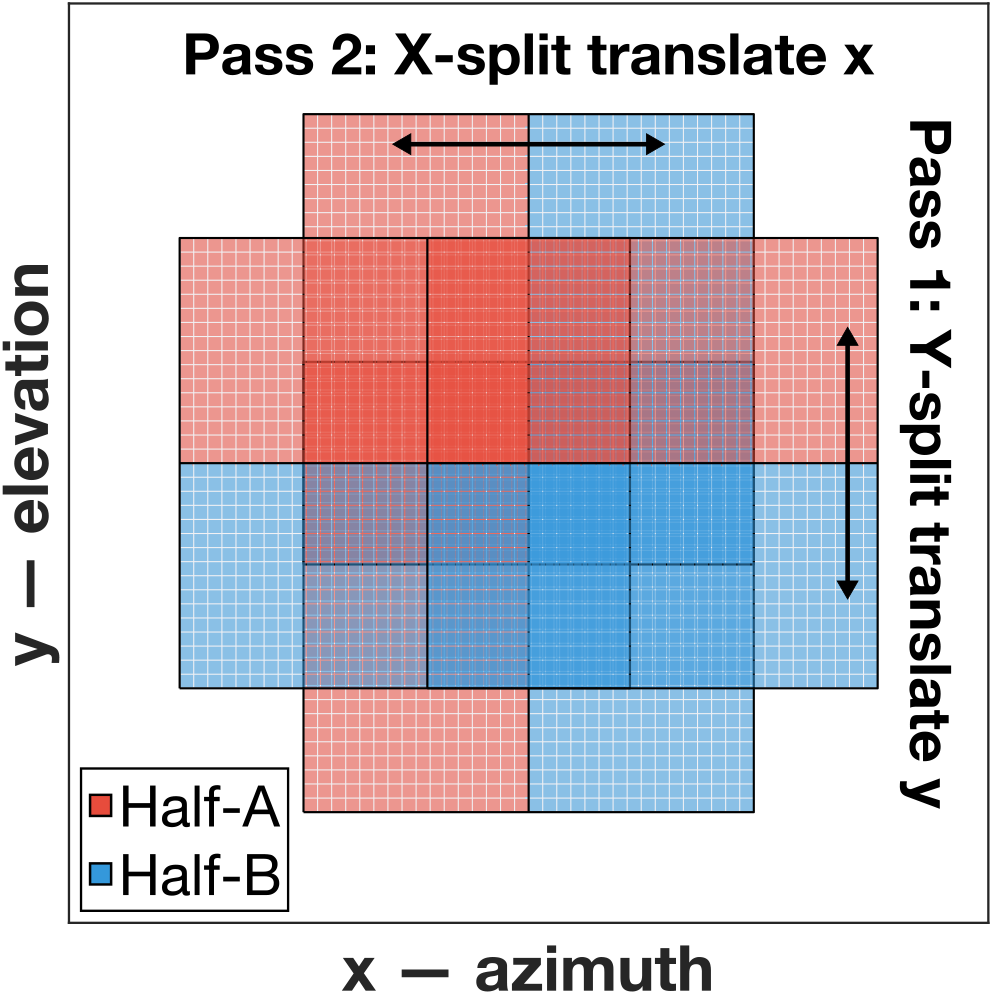
Proposed two-direction translation sweep on a monolithic matrix array (no inter-panel non-emissive regions) (Half-*A* red, Half-*B* blue). Pass 1: the Y-split (elevation top–bottom bisection) is translated along elevation (*y*). Pass 2: the X-split (azimuthal left–right bisection) is translated along azimuth (*x*).

With the full lateral-elevation plane accessible in this way, a direct *in vivo* application is longitudinal monitoring of intracranially implanted glioblastoma (GBM) [36] through the intact rodent skull. Tumors could be labeled either by intravenously injected GVs, which can extravasate through the leaky tumor vasculature [37], [38], or by engineering the GBM cells to express ARG-encoded GVs [7], [8], and in either case imaged non-invasively over days to weeks without a craniotomy. Microbubbles administered in the same session would add co-registered volumetric imaging of the tumor vasculature.

### G. Limitations

First, all image-quality measurements in Sec. III were made on a tissue-mimicking agarose phantom rather than *in vivo* (the transmit-amplitude ramp of Sec. II-C used a separate scatterer-free phantom). Second, the transcranial experiments used a single *ex vivo* mouse skull, so they do not capture inter-subject variability in cranial thickness and porosity, and the accompanying simulation used the one-way, forward-only ASM approximation of Sec. II-F. Third, the quantitative partition analysis is specific to the Vermon 32*×*32 aperture (element pitch *d* = 300 *µ*m with three *≈*330 *µ*m inter-panel non-emissive regions) at *f*_0_ = 7.81 MHz, and the best partition on monolithic or finer-pitch matrix arrays may differ. Fourth, the GV concentration and inclusion geometry were chosen to give a strong, easily quantified VxAM response, and performance at the dilute concentrations of cellular reporter-gene expression has not yet been characterized. Fifth, the resolution values of Sec. III-F are upper bounds derived from an extended inclusion boundary rather than from a point target, and the elevation field of view was not bounded by these acquisitions, because the inclusion spanned nearly the entire sweep. Sixth, the protocol records the xAM residual only after a live imaging loop has been running, so no dataset contains a first-exposure frame. The residual is stable to within 0.2 dB from the first recorded frame onward, but the fraction of the GV population lost on first insonation at the operating amplitude was not measured.

## V. Conclusion

We have demonstrated volumetric cross-amplitude modulation imaging on a 1024-element Vermon matrix array driven by four synchronized Verasonics Vantage 256 scanners. Angular spectrum simulations selected the Y-split, whose bisector is aligned with the inter-panel non-emissive regions, and a 6° steering angle, which balances peak intersection pressure against separation of the main beam from its side lobes. In a GV-loaded tissue-mimicking phantom the CTR was *≈*29 dB for single frames and *≈*49 dB with 100-frame averaging, and through an *ex vivo* mouse skull the GV well remained resolvable with a CNR of *≈*8–12 dB, against *≈*18 dB without the skull. Volumes were acquired by robotic translation, because a purely electronic asymmetric-angle sweep stayed coherent over only the central half of the aperture. Linear-field simulations of the bisecting beam thus served as surrogate predictors for designing partitions on a paneled matrix array, and their predicted trends co-varied with the measured contrast across steering angle. Through a micro-CT mouse skull model, however, one-way simulation predicted only a 1.27 dB loss in peak intersection pressure, with 20% broadening and a 0.10 mm deflection of the bisector, far less than the 12.7 dB measured drop in residual amplitude, so modeling the return path, the buckling-active volume, and the overlying soft tissue is needed to predict transcranial performance. These results are a first step toward transcranial, volumetric molecular ultrasound imaging of GVs, to be evaluated next in more skulls and *in vivo*.

## Acknowledgment

The authors thank Dr. Marie Hemelt of the University of North Carolina at Chapel Hill, Chapel Hill, NC, USA, for the animal dissection and brain extraction that provided the *ex vivo* mouse skull specimen used in the phantom measurements.

Anthropic Claude (Claude Opus 4.6, 4.7, 4.8, 5, and 5.5) was used as a coding and writing assistant throughout this work. In the software, it was used to develop and modify the Verasonics VxAM transmit sequences and the synchronized multi-Vantage acquisition and hydrophone calibration scripts, the beamforming and robotic-translation stitching pipeline, the angular spectrum scripts that simulate the partitioned transmit fields with and without the skull, and the analysis and figure-generation scripts that compute the reported CTR, CNR, and resolution values and render the figures. The angular spectrum solver, the beamformer, and the acquisition sequences on which that work builds were originally written by the authors. In the manuscript, it was used for copyediting and language polishing of author-written prose, for drafting the figure captions from panel content specified by the authors, for restructuring the LaTeX source, and for drafting bibliography entries for references selected by the authors. No experimental data was generated by the AI system, and every reported value was computed from the measured acquisitions or from the simulations described in Sec. II. The authors reviewed and verified all AI-assisted text, code, and bibliography entries, and take full responsibility for the content of the article and for the results it reports.

## Data Availability

The data that support the findings of this study are available from the corresponding author upon reasonable request. The micro-CT volume used for the aberrating-layer simulations (Sec. II-F) is a third-party dataset made publicly available by the SCRI Bone and Skeletal Mechanics Laboratory, University of Washington, and is not redistributed here.

