## Supplementary material for "Volumetric Cross-Amplitude Modulation (VxAM) for Transcranial Gas Vesicle Imaging": Supp

### Transmit Calibration and Beam Characterization

Figure S1 reports the hydrophone calibration of the drive-voltage-to-pressure mapping used throughout the manuscript, together with beam maps of the Y-split field at the operating angle. All measurements were made in water with the probe in the configuration described in Sec. II-C.

### Transmit-Amplitude Sweep

The two figures below report the transmit-amplitude sweep described in Sec. II-C of the manuscript, reconstructed two ways from the same acquisition. Figure S2 restricts the reconstruction to the bisector slab; Fig. S3 reconstructs the full transducer aperture. The comparison isolates the contribution of off-bisector gas vesicles brought above the buckling threshold by the X-wave side lobes, and is discussed in Sec. IV-C.

Both sweeps were acquired at a fixed probe position on a single scatterer-free agarose phantom, and are a separate acquisition from the contrast-versus-angle measurements of Fig. 4 in the manuscript. The contrast ratio plotted here uses the annular noise region of interest as its denominator, and is therefore not the same quantity as the contrast-to-tissue ratio (CTR) of Sec. II-K, which uses a separate tissue region of interest. Values should not be compared directly between the two.

### Volumetric Resolution

Figure S4 reports the edge-spread measurement of volumetric resolution summarized in Sec. III-F of the manuscript. No point target was available, so resolution is measured from the extended gas-vesicle inclusion boundary and every half-width reported is an upper bound.

### Skull Model Construction

This note expands on the aberrating-layer model of Sec. II-F of the manuscript. The micro-CT release identifies the specimen only as an adult *Mus musculus* dry skull, with no strain, sex, or age, and no strain is claimed here. The endocranial cavity volume recovered from the scan was ≈450 mm$^3$.

**Alignment and cropping.** The block-maximum-decimated volume was rigidly aligned by principal-component analysis of the cortical-bone voxels (counts >25 000; $1.26 \times 10^6$ voxels), which mapped the anteroposterior, left–right, and dorsoventral axes onto the grid axes with 0.48° residual misalignment, with the dorsal calvarium facing the aperture. It was cropped to the bone bounding box plus a 1 mm margin (≈25.8×14.4×12.7 mm), and the mapped volume was resampled onto the simulation grid by linear interpolation. The beam was centered laterally on the endocranial cavity, since the rostral snout displaces the centroid of the bone bounding box anteriorly.

**Intensity anchors.** Both anchors of the intensity-to-property ramp come from the specimen histogram. The lower anchor marks the separation between water and soft tissue, and the upper anchor is the cortical-bone mode. Anchoring to the mode instead of the global maximum makes the mapping insensitive to the detector- and reconstruction-dependent absolute count scale and leaves no free parameters once the histogram is fixed. Absolute radiodensity,

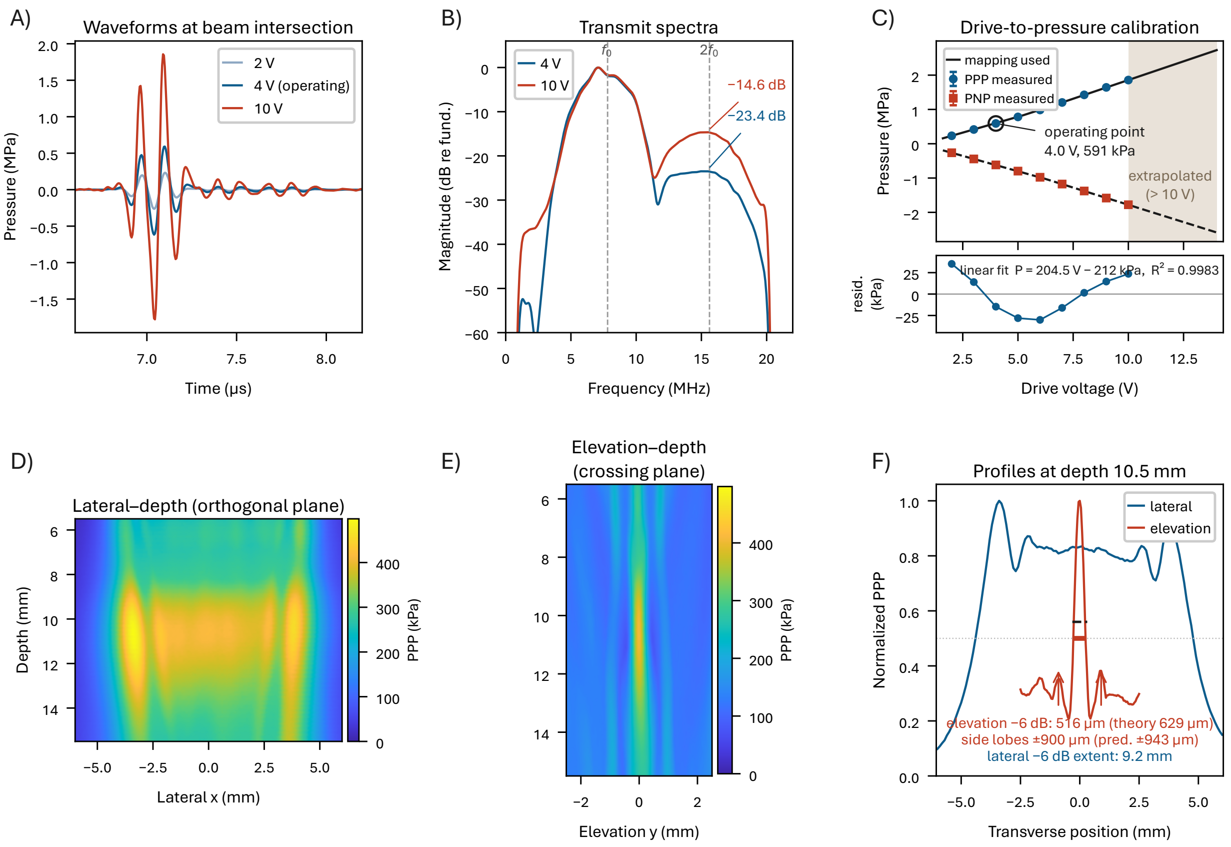


Figure S1: Hydrophone calibration and beam characterization. All measurements with an Onda HNC-0200 needle hydrophone (s/n 1177) and AH-2010 preamplifier (s/n 1477) in right-angle configuration, converted from voltage to pressure by FFT deconvolution over the 1–20 MHz factory calibration band. **(A)** Emitted pressure waveforms at 2, 4 and 10 V at the beam intersection, each the mean of ten A-lines. **(B)** Corresponding magnitude spectra at 4 and 10 V, normalized to each fundamental, with $f_0$ and $2f_0$ marked. Second-harmonic content rises from $-23.4$ dB at the 4 V operating point to $-14.6$ dB at 10 V. The received spectral peak lies near 7.1 MHz, below the nominal 7.81 MHz; the short burst's bandwidth causes this downshift, and the drive frequency is unchanged. **(C)** Peak-positive and peak-negative pressure versus drive, with the piecewise-linear voltage-to-pressure mapping used throughout this work overlaid, and residuals from a single global straight line shown below. The relationship is measurably nonlinear. A global fit ($P = 204.5\,V - 212$ kPa, $R^2 = 0.9983$) leaves systematic residuals of $\pm 35$ kPa, far exceeding the $\leq 6$ kPa repeat standard deviation, so the piecewise mapping is used instead. Shading marks the extrapolated region outside 2–10 V. **(D, E)** Peak-positive pressure maps at 3 V, $\theta = 6°$ steering (half-)angle, in the lateral–depth (orthogonal) and elevation–depth (crossing) planes on a shared scale, showing that the field is a thin sheet. Both are single-plane maps at 3 V, neither a projection nor separately normalized. **(F)** Transverse profiles through the calibration plane at 10.5 mm with $-6$ dB widths: elevation 516 $\mu$m against 629 $\mu$m predicted, side lobes at $\pm 900$ $\mu$m against $\pm 943$ $\mu$m predicted, and a 9.2 mm lateral $-6$ dB extent. Agreement of the side-lobe spacing with prediction to within $\approx 5\%$ confirms the crossing geometry. Note that (F) is taken at the 10.5 mm calibration plane. At the 11.8 mm analysis plane used elsewhere, the elevation $-6$ dB width is 616 $\mu$m against 628.8 $\mu$m predicted and the side lobes lie at $\pm 950$ $\mu$m against $\pm 943$ $\mu$m predicted, agreeing to 2% and 0.7% respectively. The measured field peaks at 10.70 mm versus 11.33 mm in simulation.

however, cannot be recovered from an uncalibrated scan, and the release contains no transmission measurement for this specimen. All comparisons therefore hold the attenuation model fixed across conditions.

**Attenuation endpoint.** At $f_0$ the upper endpoint of 8 dB cm$^{-1}$ MHz$^{-1}$ gives 62.5 dB cm$^{-1}$. It is the cortical-bone absorption of the ITRUSST transcranial simulation benchmark (4 dB cm$^{-1}$ at 500 kHz; Aubry *et al.*, 2022), which we scaled linearly with frequency; the benchmark itself prescribes no frequency law. The one-way model includes

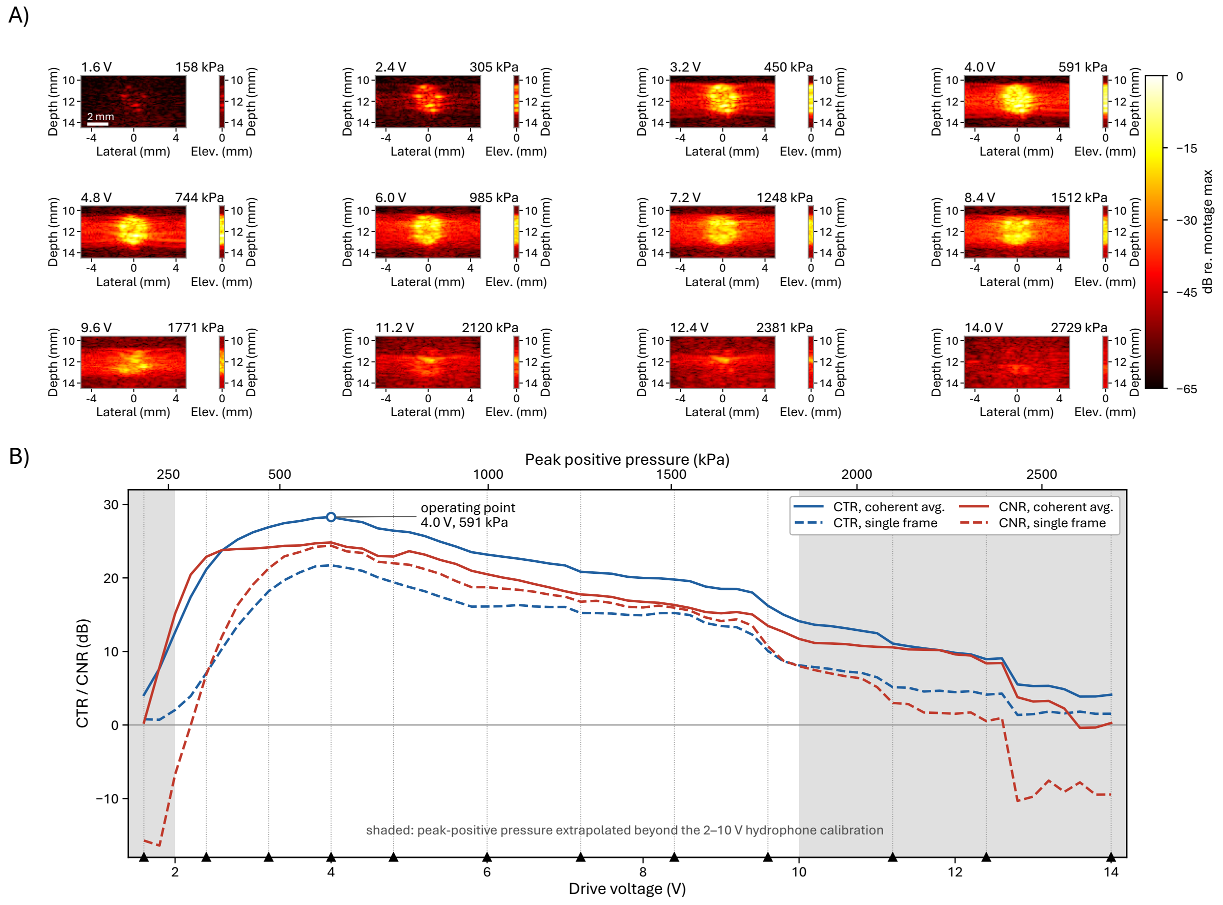


Figure S2: Contrast versus transmit pressure, bisecting-slab reconstruction. Cross-amplitude modulation (xAM) at 7.81 MHz with a Y-split aperture at a crossing half-angle of 6°, one-cycle transmit, on a gas-vesicle-loaded agarose phantom containing a 3 mm nominal cylindrical inclusion. The probe was held fixed while drive voltage was stepped from 1.6 to 14.0 V in 0.2 V increments (63 levels, 100 frames each). **(A)** Reconstructed `_XAM_slab` volumes at twelve representative levels, each shown as a lateral–depth maximum-intensity projection (left) paired with the axial–elevation projection of the same acquisition (right). All twenty-four images share one color scale and one display range, $-65$ to 0 dB relative to the brightest voxel of the montage. Per-tile normalization is deliberately not used, as it would rescale away the growth in the region exceeding threshold. Images are the 100-frame coherent (complex) average, cropped in depth to 9.5–14.5 mm. All panels are drawn at true aspect, so the elevation view is a 0.49 mm ribbon against a 9.96 mm lateral field. Scale bar 2 mm. **(B)** Contrast ratio (labeled CTR in the figure) and contrast-to-noise ratio (CNR) versus drive, for a single frame and for the 100-frame coherent average. Here $\mathrm{CTR} = 20\log_{10}(\mu_{\mathrm{GV}}/\mu_{\mathrm{noise}})$ using a concentric annulus as the denominator, and $\mathrm{CNR} = 20\log_{10}(|\mu_{\mathrm{GV}} - \mu_{\mathrm{noise}}|/\sigma_{\mathrm{noise}})$. Markers below the axis indicate the twelve levels shown in (A). The upper axis gives peak-positive pressure through the hydrophone calibration of Fig. S1; shaded regions mark drives where that pressure is extrapolated beyond the 2–10 V measured range. Contrast peaks at 28.3 dB at 4.0 V (591 kPa), the operating point used throughout, and declines gradually and monotonically above it. Single-frame CNR reaches 0 dB near 2400 kPa. The region exceeding the buckling threshold continues to expand past the contrast maximum. The area above $-30$ dB peaks at 8.4 V, and the lateral extent of that region grows from 4.4 mm at 4.0 V to 7.4 mm at 8.8 V, against a 3 mm inclusion.

refraction but not reflection, reverberation, scattering, or mode conversion, so in the model this coefficient stands in for the total transmission loss. It lies between the longitudinal absorption of cortical bone ($\approx$2.7 dB cm$^{-1}$) and the total attenuation ($\approx$13.3 dB cm$^{-1}$, measured across $\approx$8 mm human skull samples) reported by Pinton *et al.* (2012).

**Size of the simulated loss.** Two points bear on the 1.27 dB simulated drop of Sec. III-D. First, because $\alpha_0$ ramps

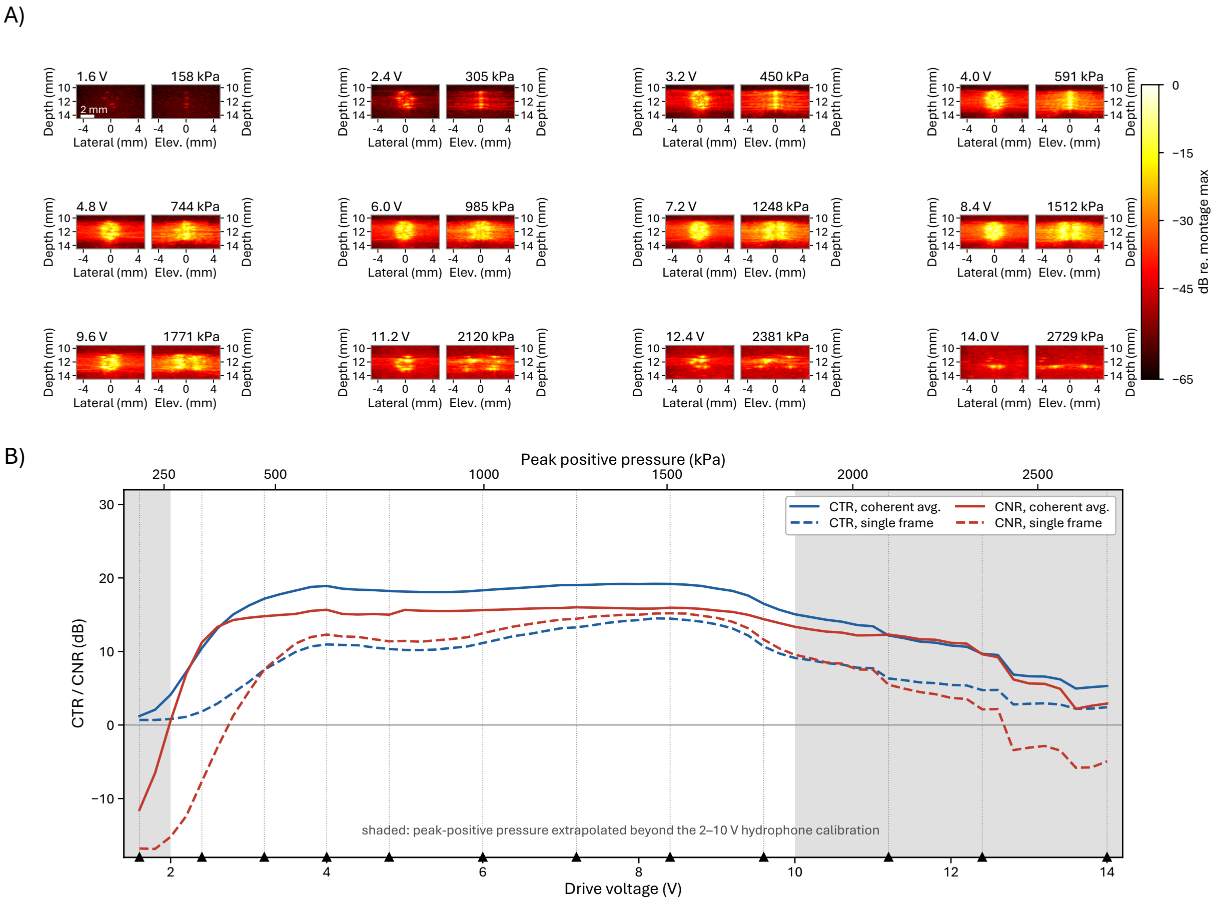


Figure S3: Contrast versus transmit pressure, volumetric reconstruction. As Fig. S2, from the same acquisition reconstructed as _XAM_vol (102 elevation planes spanning ±5 mm, versus six planes over 0.49 mm for the bisecting slab). **(A)** The elevation view is therefore a full ±5 mm image at true aspect, resolving how signal spreads in elevation as drive increases. Confinement to the bisector at low drive gives way to broad elevation coverage by 8.4 V. Display conventions are identical to Fig. S2(A). **(B)** CTR and CNR versus drive for the volumetric region of interest. No operating point is marked. The CTR maximum here falls at 8.2 V (1469 kPa), well above the 4.0 V operating point, because the volumetric region of interest spans the full ±5 mm elevation extent and continues to recruit newly super-threshold signal as drive rises, whereas the slab region of interest remains on the bisector, where the response saturates and then declines. This panel characterizes elevation behavior and should not be used to define the operating point; see Fig. S2(B).

with intensity, most calvarial voxels lie below the upper endpoint, and the path-integrated attenuation is smaller than the endpoint alone would suggest. Over a 0.3–0.4 mm calvarium the drop corresponds to an effective $\alpha_0$ of ≈4.1–5.4 dB cm$^{-1}$ MHz$^{-1}$, and to less insofar as part of the drop is refractive. Second, the phase and amplitude screens carry no impedance term. With characteristic impedances of ≈4.7 and ≈1.5 MRayl for cortical bone and water, the normal-incidence transmission loss would be ≈1.3 dB per interface, so the simulated attenuation is a lower bound on the transmit-side loss through a real calvarium.

**Aperture orientation.** With the array rotated so that its elevation (panel-stacking) direction runs anteroposteriorly, the aperture is anisotropic, with ≈5.4 mm half-aperture separation along its long axis (Sec. IV-A of the manuscript).

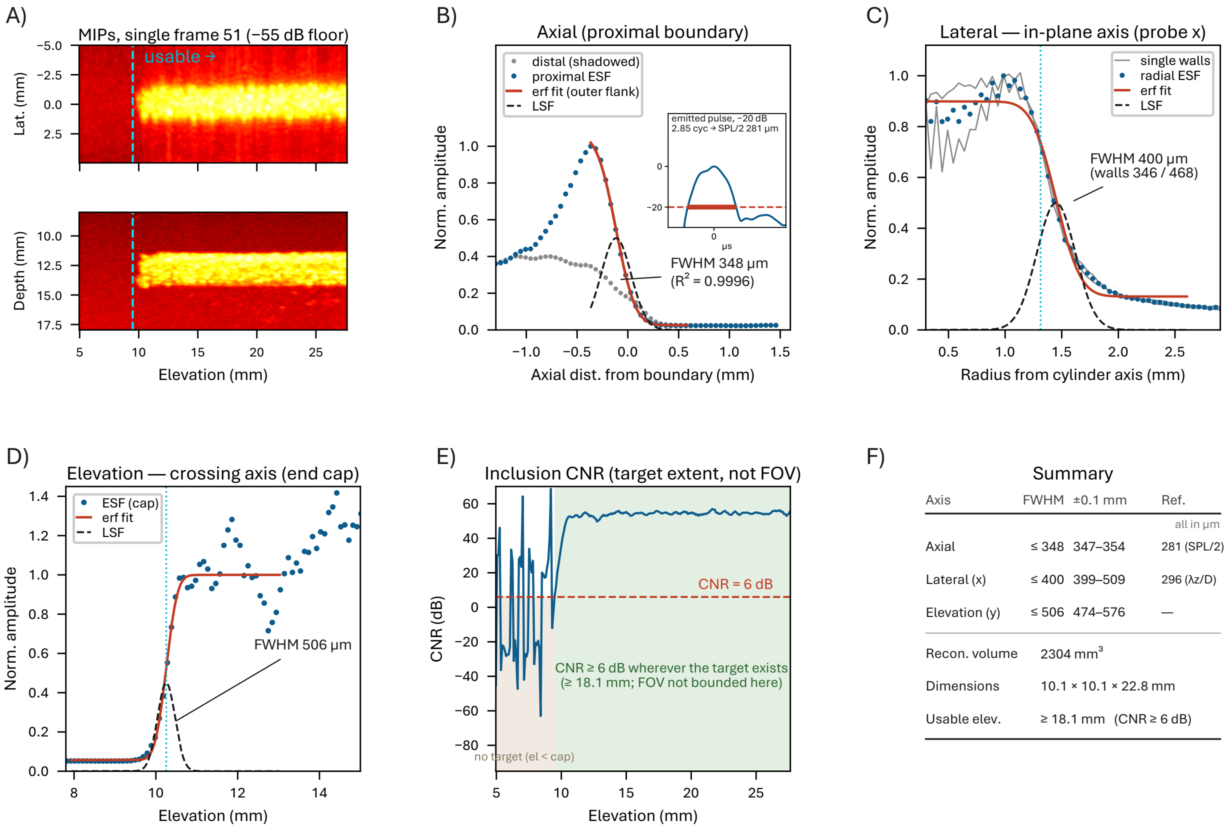


Figure S4: Volumetric resolution and target extent. Measured on the stitched no-skull phantom volume (150 robot positions at 0.2 mm pitch, of which 116 fall inside the reconstruction grid, giving a 22.8 mm elevation extent; 200 xAM frames per position, 98.6 $\mu$m isotropic grid. The contrast measurements of the manuscript use the separate 93-position, 18.1 mm sweep of Table I). No wire or bead phantom was available for this acquisition, so resolution was measured by edge-spread analysis of the 3 mm nominal cylindrical gas-vesicle inclusion, whose axis lies along elevation and which therefore presents three mutually orthogonal boundaries. The inclusion cross-section was located by a circle fitted to the 50% contour of the mean cross-section over elevations 12–26 mm. Every edge-spread function was sampled normal to that boundary, averaged over the same elevation span, fitted with an error function in a fixed window anchored on its own 50% crossing, and differentiated analytically; the full width at half maximum of the resulting line-spread function is reported. **(A)** Maximum-intensity projections, single frame 51, $-55$ dB floor; the marker indicates where the inclusion becomes detectable. **(B–E)** use the 200-frame average. **(B)** Axial, from the proximal surface, fitted on the outer flank only so as to exclude the specular interface rim. The distal surface (grey) lies in the inclusion's acoustic shadow and is not used for resolution. Inset: the measured emitted pressure pulse, 2.85 cycles at $-20$ dB. **(C)** Lateral, from both cylinder walls sampled radially within $\pm 20°$ sectors about horizontal, the only sectors in which the bright upper crescent and its acoustic shadow do not corrupt the edge. Grey: each wall alone (346 and 468 $\mu$m). **(D)** Elevation, from the distal end cap, averaged over a disk of 0.7 times the fitted radius. **(E)** Inclusion contrast-to-noise ratio versus elevation: the mean of that disk minus the mean of the same disk in the target-free slices before the end cap, divided by their standard deviation. This measures the extent over which the target is detectable; the field of view cannot be measured here, because the inclusion begins at its end cap and runs off the far edge of the reconstruction. **(F)** Summary. All FWHM values are upper bounds, the object boundary having finite width and curvature within the sampling window, both of which inflate the measured edge. The $\pm 0.1$ mm column gives the range of each FWHM when the fitted circle center is displaced by $\pm 0.1$ mm in lateral and depth. The axial reference is half the spatial pulse length of the measured emitted pulse (B). The one-cycle electrical drive is rung out to approximately three cycles by the transducer's impulse response, and it is the emitted pulse that sets axial resolution. The lateral reference is $\lambda z/D$ for the receive aperture: the beamformer uses elements within a radius $z/F_{\#}$ of each voxel at $F_{\#} = 3$ applied to the radius, so $D = 2z/3$ (effective $F_{\#} = 1.5$). Reconstruction grid 98.6 $\mu$m. Axes: lateral is the in-plane direction, parallel to the bisector plane (probe $x$); elevation is the crossing axis (probe $y$). The fitted cylinder diameter is 2.63 mm against a 3.0 mm nominal value. These data do not distinguish an undersized inclusion from an inward-biased edge criterion.
